# An emerging SARS-CoV-2 mutant evading cellular immunity and increasing viral infectivity

**DOI:** 10.1101/2021.04.02.438288

**Authors:** Chihiro Motozono, Mako Toyoda, Jiri Zahradnik, Terumasa Ikeda, Akatsuki Saito, Toong Seng Tan, Isaac Ngare, Hesham Nasser, Izumi Kimura, Keiya Uriu, Yusuke Kosugi, Shiho Torii, Akiko Yonekawa, Nobuyuki Shimono, Yoji Nagasaki, Rumi Minami, Takashi Toya, Noritaka Sekiya, Takasuke Fukuhara, Yoshiharu Matsuura, Gideon Schreiber, The Genotype to Phenotype Japan (G2P-Japan) consortium, So Nakagawa, Takamasa Ueno, Kei Sato

## Abstract

During the current SARS-CoV-2 pandemic that is devastating the modern societies worldwide, many variants that naturally acquire multiple mutations have emerged. Emerging mutations can affect viral properties such as infectivity and immune resistance. Although the sensitivity of naturally occurring SARS-CoV-2 variants to humoral immunity has recently been investigated, that to human leukocyte antigen (HLA)-restricted cellular immunity remains unaddressed. Here we demonstrate that two recently emerging mutants in the receptor binding domain of the SARS-CoV-2 spike protein, L452R (in B.1.427/429) and Y453F (in B.1.298), can escape from the HLA-24-restricted cellular immunity. These mutations reinforce the affinity to viral receptor ACE2, and notably, the L452R mutation increases protein stability, viral infectivity, and potentially promotes viral replication. Our data suggest that the HLA-restricted cellular immunity potentially affects the evolution of viral phenotypes, and the escape from cellular immunity can be a further threat of the SARS-CoV-2 pandemic.

**Graphical Abstract:** 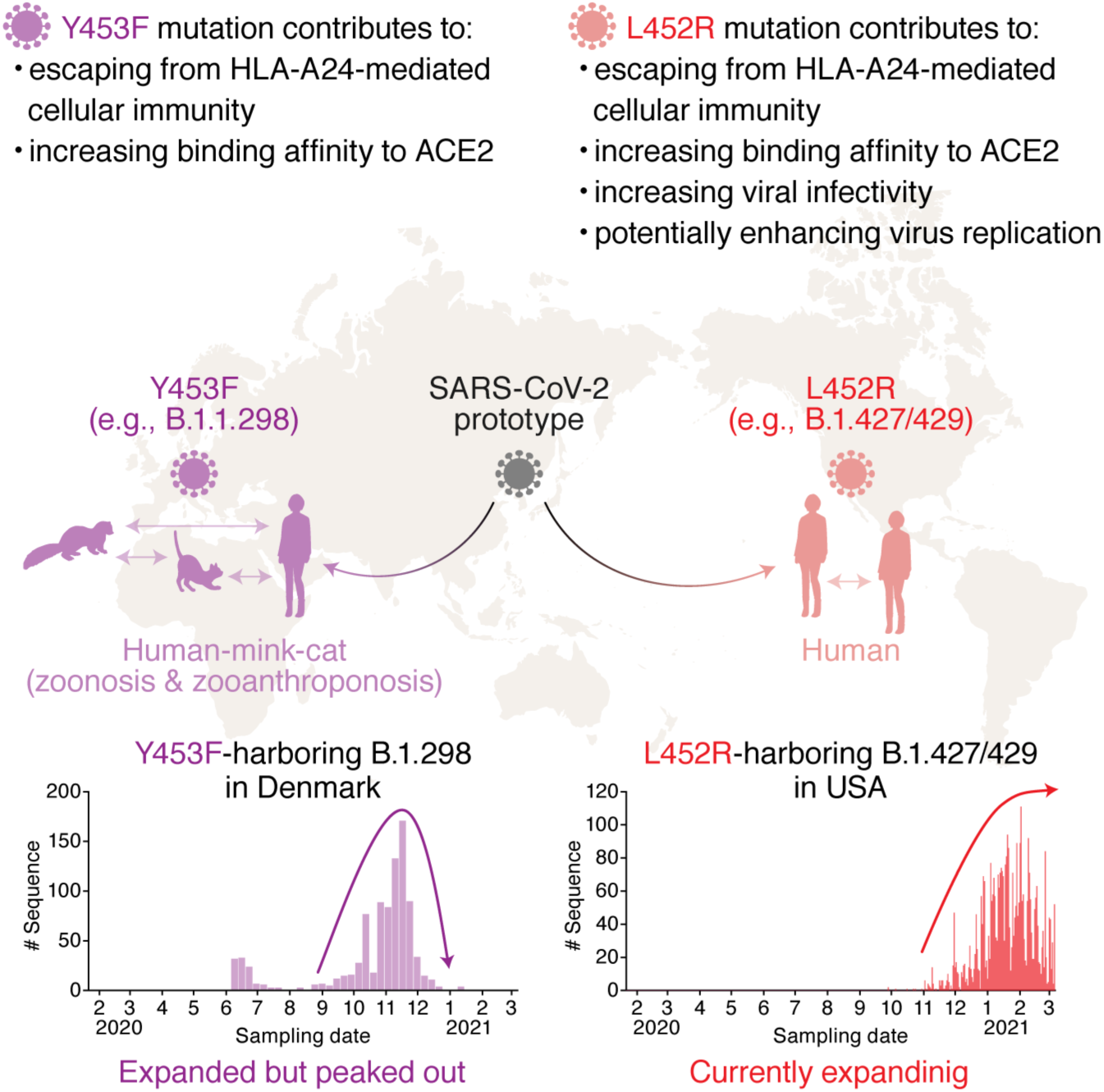

## Introduction

Coronavirus disease 2019 (COVID-19) is an infectious disease caused by severe acute respiratory syndrome coronavirus 2 (SARS-CoV-2). Since an unusual outbreak in Wuhan, Hubei province, China, in December 2019 (Wu et al., 2020; Zhou et al., 2020), SARS-CoV-2 has rapidly spread all over the world, and as of March 2021, SARS-CoV-2 is an ongoing pandemic: more than one hundred million cases of infections have been reported worldwide, and more than two million people died of COVID-19 (WHO, 2020a).

During the current pandemic, a variety of SARS-CoV-2 mutants have emerged and some of them have dominantly spread [reviewed in (Plante et al., 2021)]. A well-studied SARS-CoV-2 mutant harbors D614G substitution in the spike (S) protein. Recent studies have revealed that the D614G mutation increases the SARS-CoV-2 binding affinity to ACE2, the SARS-CoV-2 receptor (Ozono et al., 2021; Yurkovetskiy et al., 2020; Zhou et al., 2021), infectivity (Ozono et al., 2021; Yurkovetskiy et al., 2020; Zhou et al., 2021), fitness (Hou et al., 2020; Plante et al., 2020; Zhou et al., 2021), and transmissibility in the human population (Volz et al., 2021). However, there is no evidence suggesting that the D614G variant is associated with viral pathogenicity and lethality (Hou et al., 2020; Korber et al., 2020; Plante et al., 2020). Additionally, since the fall of 2020, new SARS-CoV-2 variants such as B.1.1.7 (also known as a variant of concern 202012/01 or 20I/501Y.V1), B.1.351 (also known as 20H/501Y.V2), and P.1 (also known as 501Y.V3) lineages emerged in the UK, South Africa, and Brazil, respectively, and have rapidly spread worldwide (CDC, 2020). At the end of 2020, another lineage, B.1.427/429 (also known as CAL.20C), has been predominant particularly in California state, the USA (Deng et al., 2021; Zhang et al., 2021). Moreover, cross-species viral infection can accelerate the emergence of diversified viruses [reviewed in (Banerjee et al., 2021; Parrish et al., 2008)]. In the case of SARS-CoV-2, a variety of mammals such as nonhuman primates (Chandrashekar et al., 2020; Munster et al., 2020; Yu et al., 2020) and carnivores (Halfmann et al., 2020; Kim et al., 2020; Shi et al., 2020) are prone to its infection (Damas et al., 2020; Martinez-Hernandez et al., 2020; OIE, 2021). Strikingly, the emergence of a SARS-CoV-2 variant, B.1.298, has likely to be associated with the outbreak in farmed minks in Denmark (Koopmans, 2021; WHO, 2020b), and phylogenetic analysis has provided evidence of mink-to-human transmission of SARS-CoV-2 within Danish mink farms (Oude Munnink et al., 2021). Because newly emerging variants can potentially change viral infectivity, transmissibility and pathogenicity, deep monitoring of the SARS-CoV-2 strains circulating globally and locally and evaluating the effects of mutations detected on virological characteristics are urgent and crucial.

The emergence of mutated viruses is mainly due to error-prone viral replication, and the spread of emerged variants is attributed to the escape from immune selective pressures [reviewed in (Duffy et al., 2008)]. In fact, several SARS-CoV-2 mutants can be resistant to the neutralization mediated by the antibodies from COVID-19 patients (Baum et al., 2020; Chen et al., 2021; Liu et al., 2021c; McCarthy et al., 2021; Weisblum et al., 2020) as well as those from vaccinated individuals (Liu et al., 2021b). Although the B1.1.7 variant is sensitive to convalescent and vaccinated sera (Collier et al., 2021; Garcia-Beltran et al., 2021; Shen et al., 2021; Supasa et al., 2021; Wang et al., 2021), the B.1.351 and P.1 variants are relatively resistant to anti-SARS-CoV-2 humoral immunity (Garcia-Beltran et al., 2021; Hoffmann et al., 2021a; Wang et al., 2021).

In addition to the humoral immunity mediated by neutralizing antibodies, another protection system against pathogens is the cellular immunity mediated by cytotoxic T lymphocytes (CTLs) [reviewed in (Fryer et al., 2012; Leslie et al., 2004)]. CTLs recognize the nonself epitopes that are presented on virus-infected cells via human leukocyte antigen (HLA) class I molecules, and therefore, the CTL-mediated antiviral immunity is HLA-restricted [reviewed in (La Gruta et al., 2018)]. Recent studies have reported the HLA-restricted SARS-CoV-2-derived epitopes that can be recognized by human CTLs (Kared et al., 2021; Kiyotani et al., 2020; Nelde et al., 2021; Schulien et al., 2021; Wilson et al., 2021). More importantly, Bert et al. have recently reported that the functionality of virus-specific cellular immunity is inversely correlated to the COVID-19 severity (Le Bert et al., 2021). Therefore, it is conceivable to assume that the HLA-restricted CTLs play crucial roles in controlling SARS-CoV-2 infection and COVID-19 disorders. However, comparing to humoral immune responses, it remains unclear whether the SARS-CoV-2 variants can potentially escape from cellular immunity.

In this study, we investigate the possibility for the emergence of the SARS-CoV-2 mutants that can escape from the HLA-restricted cellular immunity. We demonstrate that at least two naturally occurring substitutions in the receptor binding motif (RBM; residues 438-506) of the SARS-CoV-2 S protein, L452R and Y453F, which were identified in the two major variants, B.1.427/429 (L452R) and B1.1.298 (Y453F), can be resistant to the cellular immunity in the context of HLA-A*24:02, an allele of HLA-I. More intriguingly, the L452R and Y453F mutants increase the binding affinity to ACE2, and the experiments using pseudoviruses show that the L452R substitution increases viral infectivity. Furthermore, we artificially generate the SARS-CoV-2 harboring these point mutations by reverse genetics and demonstrate that the L452R mutants enhance viral replication capacity.

## Results

### Evasion from the HLA-A24-restricted CTL responses by acquiring mutations in the RBM of SARS-CoV-2 S protein

We set out to address the possibility of the emergence of the naturally occurring mutants that can potentially confer the resistance to antigen recognition by HLA-restricted cellular immunity. A bioinformatic study has suggested that the 9-mer peptide in the RBM, NYNYLYRLF (we designate this peptide “NF9”), which spans 448-456 in the S protein, can be the potential epitope presented by HLA-A24 (Kiyotani et al., 2020), an HLA-I allele widely distributed all over the world and particularly predominant in East and Southeast Asian area (**Table S1**). Additionally, three immunological analyses using COVID-19 convalescents have shown that the NF9 peptide is an immunodominant epitope presented by HLA-A*24:02 (Gao et al., 2021; Hu et al., 2020; Kared et al., 2021). To verify these observations, we obtained the peripheral blood mononuclear cells (PBMCs) from nine COVID-19 convalescents with HLA-A*24:02 and stimulated these cells with the NF9 peptide. As shown in **Figure 1A**, a fraction of CD8^+^ T cells upregulated two activation markers, CD25 and CD137, in response to the stimulation with NF9. In the nine samples of COVID-19 convalescents with HLA-A*24:02, the percentage of the CD25^+^CD137^+^ cells in the presence of the NF9 peptide (5.3% in median) was significantly higher than that in the absence of the NF9 peptide (0.49% in median) (**Figure 1B**; *P*=0.016 by Wilcoxon signed-rank test). Additionally, the stimulation with the NF9 peptide did not upregulate CD25 and CD137 in the CD8^+^ T cells of three seronegative samples with HLA-A*24:02 and the percentage of the CD25^+^CD137^+^ cells in seronegative samples (0.93% in median) was significantly lower than that in COVID-19 convalescent samples (**Figure 1B**; *P*=0.011 by Mann-Whitney U test). Consistent with previous reports (Gao et al., 2021; Hu et al., 2020; Kared et al., 2021; Kiyotani et al., 2020), our data suggest that the NF9 peptide is an immunodominant HLA-A*24:02-restricted epitope recognized by the CD8^+^ T cells of COVID-19 convalescents in our cohort.

**Figure 1.**
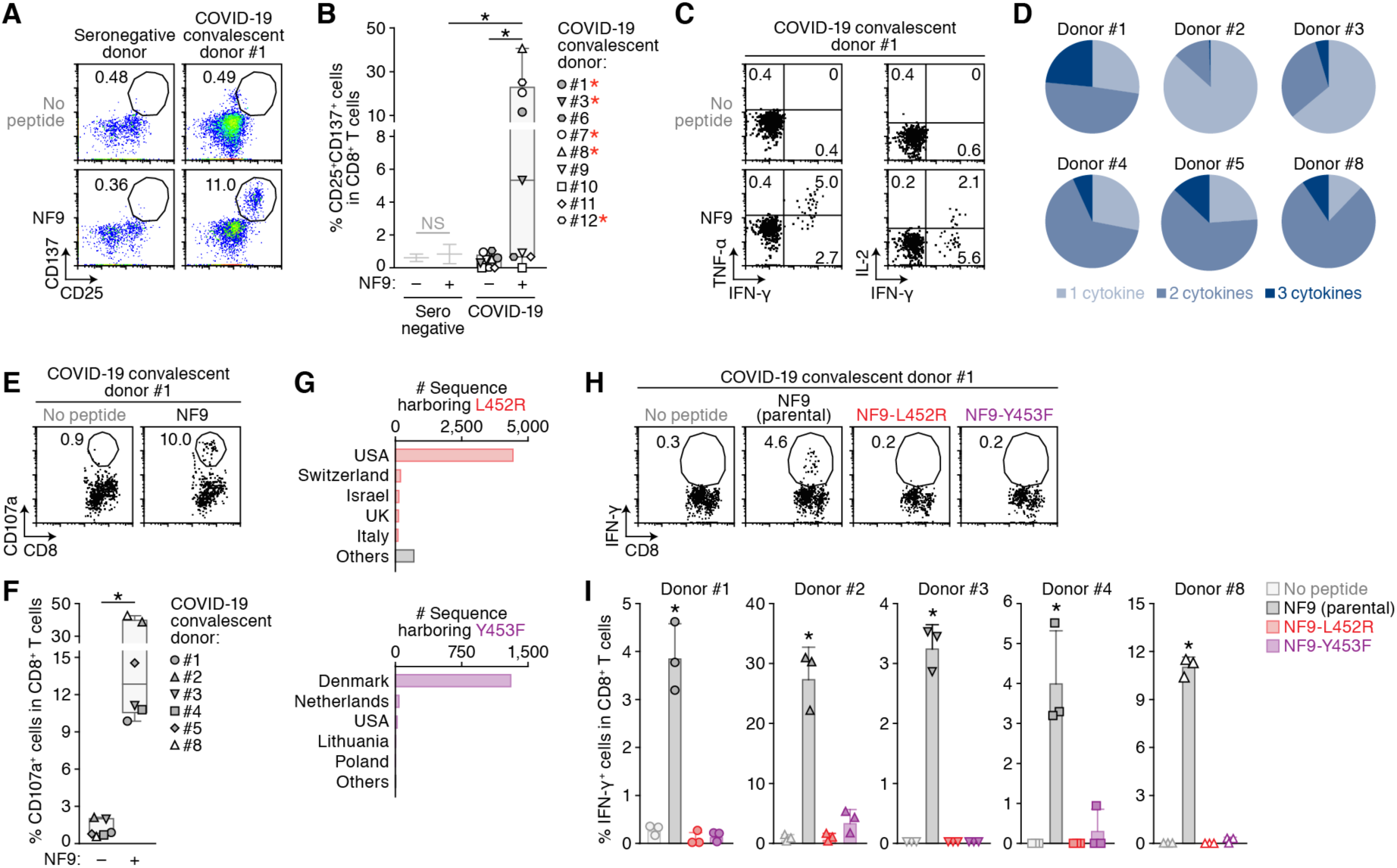
Escape of the two naturally occurring SARS-CoV-2 mutations from the S RBM-specific CD8^+^ T cells. (**A and B**) Detection of the HLA-A24-restricted NF9-specific CTLs. The HLA-A*24:02-positive CTL lines of 3 seronegative donors and 9 COVID-19 convalescents were stimulated with or without 1 μM NF9 peptide (NYNYLYRLF, residues 448-456 of the SARS-CoV-2 S protein). Representative FACS plots showing the surface expression of CD25 and CD137 in the CD8^+^ T cell subset (i.e., CD3^+^CD8^+^ cells) of a seronegative donor (left) and a COVID-19 convalescent donor #1 (right) (**A**) and the median of the percentage of CD25^+^CD137^+^ cells in CD8^+^ T cells (**B**) are shown. In **B**, the COVID-19 convalescent samples >3 SD of the median of NF9-stimulated seronegative samples are indicated with red asterisks. (**C and D**) Multifunctionality of the HLA-A24-restricted NF9-specific CTLs. The HLA-A*24:02-positive CTL lines of 6 COVID-19 convalescents were stimulated with or without 10 nM NF9 peptide. Representative FACS plots showing the intracellular expression of IFN-γ, TNF-α and IL-2 in the CD8^+^ T cell subset of a COVID-19 convalescent (donor #1) (**C**) and the pie charts showing the proportion of cytokine positive cells in each convalescent sample (**D**) are shown. (**E and F**) Potential killing activity of the HLA-A24-restricted NF9-specific CTLs. The HLA-A*24:02-positive CTL lines of 6 COVID-19 convalescents were stimulated with the C1R-A2402 cells pulsed with or without 10 nM NF9 peptide. Representative FACS plots showing the surface expression of CD107a in the CD8^+^ T cell subset of a COVID-19 convalescent (donor #1) (**E**) and the median of the percentage of CD107a^+^ cells in CD8^+^ T cells (**F**) are shown. (**G**) Distribution of the L452R and Y453F mutants during the current pandemic. The top 5 countries where the variants harboring the L452R (top) and Y453F (bottom) mutations are shown. The raw data are summarized in **Table S2**. (**H and I**) Mutations escaped from the HLA-A24-restricted NF9-specific CTLs. The HLA-A*24:02-positive CTL lines of 5 COVID-19 convalescents were stimulated with 1 nM NF9 peptide or its derivatives: NF9-L452R (NYNY<u>R</u>YRLF) and NF9-Y453F (NYNYL<u>F</u>RLF). Representative FACS plots showing the intracellular expression of IFN-γ in the CD8^+^ T cell subset of a COVID-19 convalescent (donor #1) (**H**) and the mean of the percentage of IFN-γ^+^ cells in CD8^+^ T cells (**I**) are shown. In **A, E and H**, the numbers in the FACS plot represent the percentage of gated cells in CD8^+^ T cells. In **C**, the number represents the percentage of the cells in each quadrant. In **B**, a statistically significant difference (*, *P*<0.05) between SARS-CoV-2 seronegative and COVID-19 convalescent samples is determined by Mann-Whitney U test, and a statistically significant difference (*, *P*<0.05) between with and without NF9 peptide in COVID-19 convalescent samples is determined by Wilcoxon signed-rank test. In **F**, each symbol of the COVID-19 convalescent data represents the mean of technical triplicate. Statistically significant differences (*, *P*<0.05) between with and without NF9 peptide in COVID-19 convalescent samples are determined by Wilcoxon signed-rank test. In **I**, the assay was performed in triplicate, and the means are shown with SD. Statistically significant differences (*, *P*<0.05) versus “no are determined by ANOVA with multiple comparisons by Bonferroni correction. See also **Figure S1** and **Tables S1 and S2**.

We next assessed the profile of cytokine production by the NF9 stimulation. As shown in **Figure 1C**, the stimulation with the NF9 peptide induced the production of IFN-γ, TNF-α and IL-2 in the CD8^+^ T cells of a COVID-19 convalescent. The analysis using six COVID-19 convalescent samples showed that CD8^+^ T cells produce multiple cytokines in response to the NF9 stimulation (**Figure 1D**), demonstrating the multifunctional nature of the NF9-specific CD8^+^ T cells of COVID-19 convalescents. Moreover, the cytotoxic potential of the NF9-specific CD8^+^ T cells was assessed by staining with surface CD107a, a degranulation marker (**Figure 1E**). As shown in **Figure 1F**, the percentage of CD107a^+^ cells in the CD8^+^ T cells with the NF9 peptide (12.9% in median) was significantly higher than that without the NF9 peptide (0.83% in median) (; *P*=0.031 by Wilcoxon signed-rank test), suggesting the cytotoxic potential of the NF9-specific CD8^+^ T cells.

To assess the presence of naturally occurring variants harboring mutations in this region (residues 448-456 in the S protein), we analyzed the diversity of SARS-CoV-2 during the current pandemic. We downloaded 750,243 viral genome sequences from the global initiative on sharing all influenza data (GISAID) database (https://www.gisaid.org; as of March 15, 2021). The L452R substitution was most frequent among the sequences analyzed (5,677 sequence), and 1,380 sequences reported contained the Y453F substitution (**Table 1**). Notably, the B.1.427/429 and B.1.1.298 lineages (CDC, 2020) in the PANGO lineages (https://cov-lineages.org/index.html) mainly harbor the L452R and Y453F mutations, respectively (**Figure 1G** **and Table S2**).

**Table 1.** Naturally occurring mutations in the residues 448-456 of the SARS-CoV-2 S protein

| Mutation | # Sequence | Date, first detected | Country, first detected | Dominant PANGO lineage | Date, first detected in the dominant lineage | Country, first detected in the dominant lineage | Nonhuman hosts detected <sup>a</sup> |
| --- | --- | --- | --- | --- | --- | --- | --- |
| L452R | 5,677 | March 17, 2020 | Denmark | B.1.427/429 | July 6, 2020 | Mexico | Gorilla (1) |
| Y453F | 1,380 | April 20, 2020 | Denmark | B.1.298 | April 20, 2020 | Denmark | Mink (339) and cat (3) |
| N450K | 181 | March 27, 2020 | UK | B.1.214.2 | December 5, 2020 | UK | - |
| L452M | 140 | April 13, 2020 | Japan | B.1.22/36 | July 2, 2020 | Netherlands | Mink (32) |
| L452Q | 111 | March 6, 2020 | Spain | B.1.1.374 | October 23, 2020 | Belarus | - |

<sup>a</sup> The number in parenthesis indicates the number of sequence reported.

To address the possibility that the naturally occurring mutations in the NF9 region, L452R and Y453F, evade the NF9-specific CD8^+^ T cells of HLA-A24-positive COVID-19 convalescents, two NF9 derivatives containing either L452R or Y453F substitution (NF9-L452R and NF9-Y453F) were prepared and used for the stimulation experiments. As shown in **Figure S1**, parental NF9 induced IFN-γ expression in a dose-dependent manner. In contrast, the induction level of IFN-γ expression by the NF9-Y453F derivative was significantly lower than that by parental NF9, and more intriguingly, the NF9-L452R derivative did not induce IFN-γ expression even at the highest concentration tested (10 nM) (**Figure S1**). In the five HLA-A24-positive COVID-19 convalescent samples, parental NF9 peptide significantly induced IFN-γ expression, while the NF9-L452R and NF9-Y453F derivatives did not (**Figures 1H and 1I**). Altogether, these results suggest that the NF9 peptide, which is derived from the RBM of SARS-CoV-2 S protein, is an immunodominant epitope of HLA-A24, and two naturally occurring mutants, L452R and Y453F, evade the HLA-A24-restricted cellular immunity.

### Augmentation of the binding affinity to ACE2 by the L452 and Y453 mutations

We next addressed whether the mutations of interest affect the efficacy of virus infection. Structural analyses have shown that the Y453 and N501 residues in the RBM are located on the interface between the SARS-CoV-2 RBM and human ACE2 and directly contribute to the binding to human ACE2, while the L452 residue is not on the RBM-ACE2 interface (Lan et al., 2020; Wang et al., 2020; Zhao et al., 2020) (**Figure 2A**). To directly assess the effect of these mutations in the RBM on the binding affinity to ACE2, we prepared the yeasts expressing parental SARS-CoV-2 receptor binding domain RBD (residues 336-528) and its derivatives (L452R, Y453F and N501Y) and performed *in vitro* binding assay using the yeast surface display of the RBD and soluble ACE2 protein. Consistent with recent studies including ours (Supasa et al., 2021; Zahradník et al., 2021b), the N501Y mutation, which is a common mutation in the B1.1.7, B1.351 and P.1 variants [reviewed in (Plante et al., 2021)] as well as the Y453F mutation (Bayarri-Olmos et al., 2021; Zahradník et al., 2021b) significantly increased the binding affinity to human ACE2 (**Figures 2B and 2C**; RBD parental *K*_D_ = 2.05 ± 0.26 nM; RBD N501Y *K*_D_ = 0.59 ± 0.03 nM; and RBD Y453F *K*_D_ = 0.51 ± 0.06 nM). We also found that the L452R mutant significantly increased the binding affinity to human ACE2 (**Figures 2B and 2C**; RBD L452R *K*_D_ = 1.20 ± 0.06 nM). Intriguingly, the L452R mutations increased the surface expression, which reflects protein stability (Traxlmayr and Obinger, 2012), while the Y453F and N501Y mutations decreased (**Figure 2D**).

**Figure 2.**
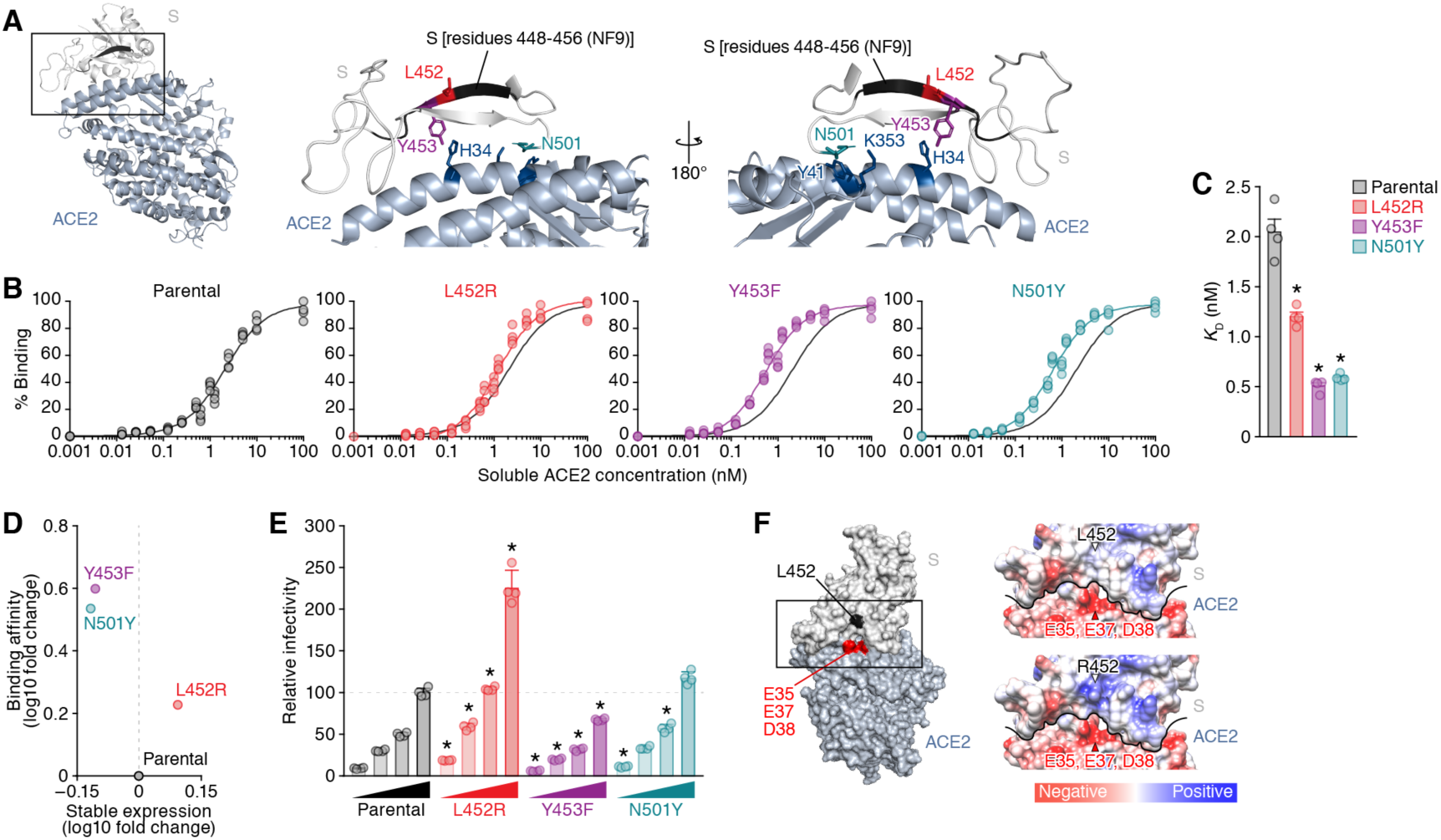
Increase of the binding affinity to ACE2 and viral infectivity by the L452 mutation. (**A**) Location of the NF9 peptide (residues 448-456) in the cocrystal structure of the SARS-CoV-2 S and human ACE2 proteins (PDB: 6M17) (Yan et al., 2020). An overview (left), the enlarged view of the boxed area in the left panel (middle) and the view of the middle panel rotated 180° on the y-axis (right) are shown. The residues 448-456 of SARS-CoV-2 S (corresponding to the NF9 peptide) are shown in black. (**B-D**) Binding affinity of SARS-CoV-2 S RBD to ACE2 by yeast surface display. The percentage of the binding of the SARS-CoV-2 S RBD expressing on yeast to soluble ACE2 (**B**) and the *K*_D_ values (**C**) are shown. (**D**) The level of stable expression of the SARS-CoV-2 RBD on yeast (x-axis) and the binding affinity to ACE2 (y-axis) compared to parental RBD. In **B**, the fitting curve of parental RBD is shown in all panels as black lines. (**E**) Pseudovirus assay. The HIV-based reporter virus pseudotyped with the parental SARS-CoV-2 S or its derivatives (L452R, Y453F and N501Y) were inoculated into the 293 cells transiently expressing human ACE2 and TMPRSS2 at 4 different doses (1, 3, 5 and 10 ng p24 antigens). The percentages of the infectivity compared to the virus pseudotyped with parental S (10 ng p24 antigen) are shown. (**F**) Gain of electrostatics complementarity by the L452R substitution. (Left) The surface structure of the SARS-CoV-2 S and ACE2 (PDB: 6M17) (Yan et al., 2020). The residue 452 of the SARS-CoV-2 S and the negatively charged patch on ACE2 (residues E35, E37 and D38) are indicated by black and red. The boxed area is enlarged in the upper right panel. (Right) Coulombic surface coloring at the structures of the SARS-CoV-2 S and ACE2 (PDB: 6M17) (Yan et al., 2020) (top) and a model of the L452R substitution (bottom). The black line indicates the border between SARS-CoV-2 S and ACE2. In **B and E**, these assays were performed in quadruplicate. In **C**, statistically significant differences (*, *P*<0.05) versus parental S are determined by Mann-Whitney U test. In **E**, statistically significant differences (*, *P*<0.05) versus parental S at the same dose are determined by ANOVA with multiple comparisons by Bonferroni correction.

### Increase of pseudovirus infectivity by the L452R mutation

To directly analyze the effect of the mutations of interest on viral infectivity, we prepared the HIV-1-based reporter virus pseudotyped with the SARS-CoV-2 S protein and its mutants and the 293 cells transiently expressing human ACE2 and TMPRSS2. As shown in **Figure 2E**, although the N501Y mutation faintly affected viral infectivity in this assay, the L452R mutations significantly increased viral infectivity compared to parental S protein. In contrast to the yeast display assay (**Figures 2B and 2C**), the infectivity of the Y453F mutant was significantly lower than that of parental S protein (**Figure 2E**). Altogether, these findings suggest that the L452R substitution increases the binding affinity of the SARS-CoV-2 RBD to human ACE2, protein stability, and viral infectivity. Although the L452 residue is not directly located at the binding interface (**Figure 2A**), structural analysis and *in silico* mutagenesis suggested that the L452R substitution can cause a gain of electrostatics complementarity (Selzer et al., 2000) (**Figure 2F**). Because the residue 452 is located close proximity to the negatively charged patch of ACE2 residues (E35, E37, D38), the increase of viral infectivity by the L452R substitution can be attributed to the increase in the electrostatic interaction with ACE2.

### Promotion of SARS-CoV-2 replication in cell cultures by the L452 mutation

To investigate the effect of the mutations in the RBM on viral replication, we artificially generated the recombinant SARS-CoV-2 viruses that harbor the mutations of interest as well as parental recombinant virus by a reverse genetics system (Torii et al., 2021). The nucleotide similarity of SARS-CoV-2 strain WK-521 (GISAID ID: EPI_ISL_408667) (Matsuyama et al., 2020), the backbone of the artificially generated recombinant SARS-CoV-2, to strain Wuhan-Hu-1 (GenBank: NC_045512.2) (Wu et al., 2020) is 99.91% (27 nucleotides difference) and the sequences encoding the S protein between these two strains are identical, indicating that the strain WK-521 is a SARS-CoV-2 prototype. We verified the insertions of the targeted mutations in the generated viruses by direct sequencing (**Figure 3A**) and performed virus replication assay using these recombinant viruses. As shown in **Figure 3B**, we revealed that the growth of the L452R mutant in VeroE6/TMPRSS2 cells was significantly higher than that of parental virus. Together with the findings in the binding assay (**Figures 2B-2D**) and the assay using pseudoviruses (**Figure 2E**), our results suggest that the L452R mutation potentially increase viral replication.

**Figure 3.**
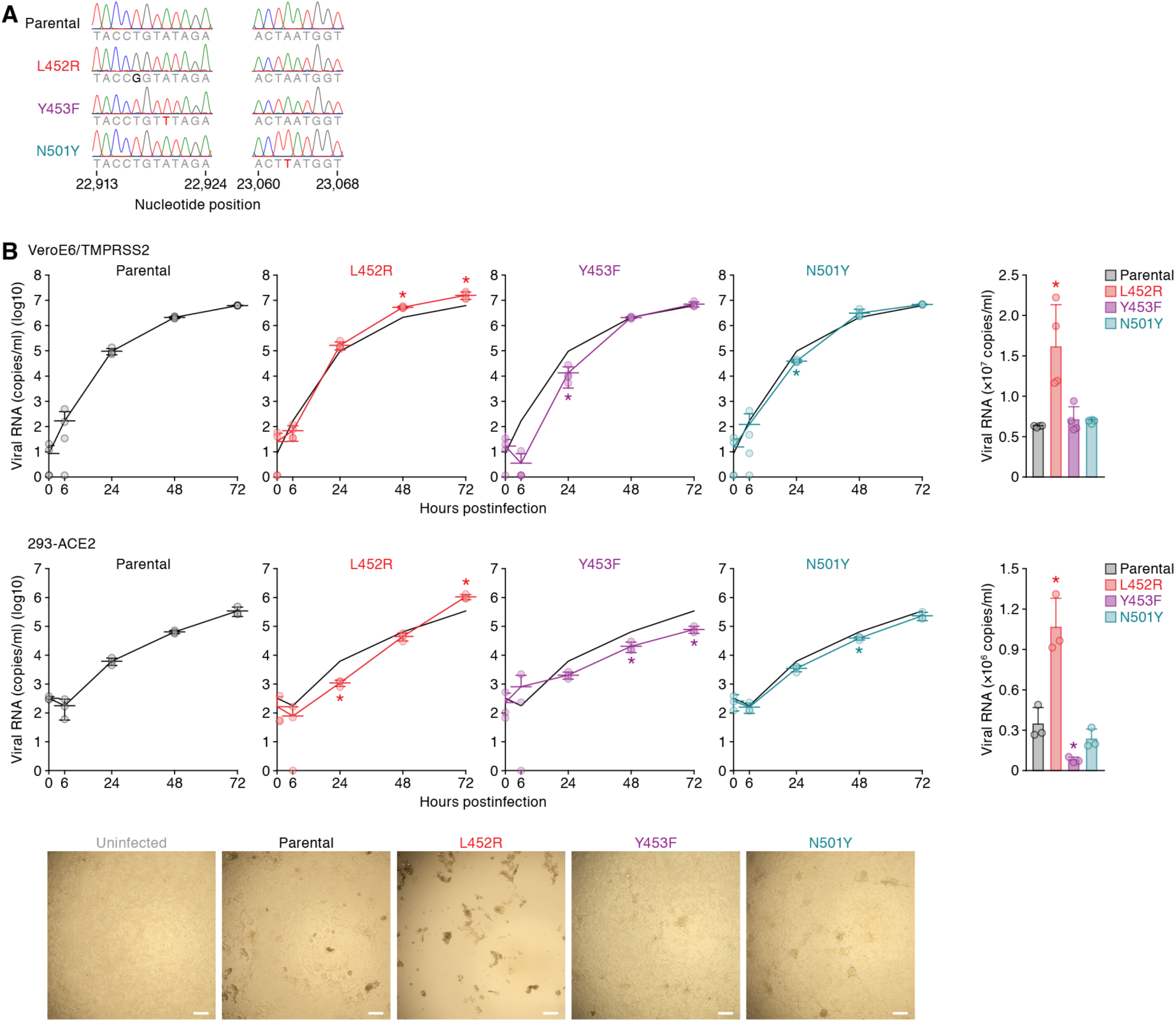
Enhancement of viral replication by the L452 mutation. (**A**) Chromatograms of the mutated regions of the SARS-CoV-2 viruses artificially generated by reverse genetics. The chromatograms of the nucleotide positions 22,913-22,924 (left) and 23,060-23,068 (right) of parental SARS-CoV-2, the L452R, Y453F and N501Y mutants are shown. (**B**) Growth kinetics of parental SARS-CoV-2 and the SARS-CoV-2 mutants. Parental SARS-CoV-2, the L452R, Y453F and N501Y mutants (100 pfu) were inoculated into VeroE6/TMPRSS2 cells (top) and 293-ACE2 cells (middle) and the copy number of viral RNA in the culture supernatant was quantified by real-time RT-PCR. (Left) The growth curve of the viruses inoculated. The result of parental virus is shown in all panels as a black line. (Right) The amount of viral RNA in the culture supernatant at 72 h postinfection. The assays were performed in quadruplicate (VeroE6/TMPRSS2 cells) or triplicate (293-ACE2 cells), and statistically significant differences (*, *P*<0.05) versus parental S are determined by Student’s *t* test. (Bottom) Representative figures of the blight fields of 293-ACE2 cells infected with the viruses indicated at 72 h post infection are also shown. Bars, 200 μm.

### Dynamics of the spread of the RBM mutants during the current pandemic

We finally assessed the epidemic dynamics of the naturally occurring variants containing the substitutions in L452 and Y453. As shown in **Figure 4A** **and Table S3**, the L452R mutants were mainly found (3,967 sequences) in the B.1.427/B.1.429 lineage that forms a single clade (Deng et al., 2021). Although the L452R mutant was first detected the B.1.39 lineage in Denmark on March 17, 2020 (GISAID ID: EPI_ISL_429311) (**Table 1**), this variant did not spread. The oldest sequence that contains the L452R mutation in the B.1.427/B.1.429 lineage was isolated in Quintana Roo state, Mexico, on July 6, 2020 (GISAID ID: EPI_ISL_942929) (**Table 1**), and the L452R-harboring mutants have been first detected in California state, the USA, on September 28, 2020 (GISAID ID: EPI_ISL_730092 and EPI_ISL_730345) (**Figure 4B**). The B.1.427/B.1.429 lineage harboring the L452R mutation has started expanding in California state, the USA, at the beginning of November, 2020 (**Figure 4B, top**). In 2021, this lineage has expanded throughout the USA, and currently, is one of the most predominant lineages in the country (**Figure 4B, bottom and Table S4**).

**Figure 4.**
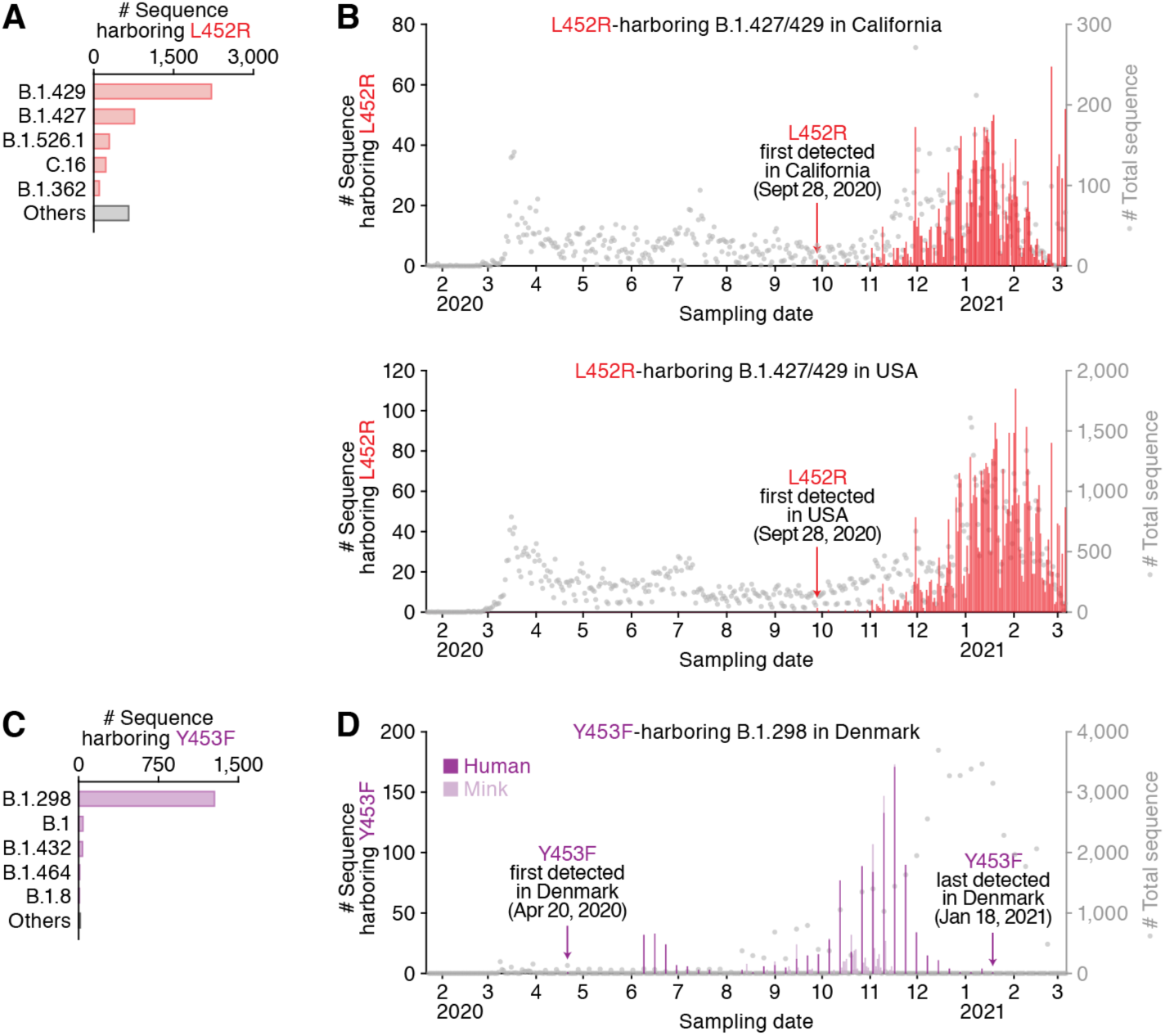
Epidemic dynamics of the B.1.298 and B.1.427/429 lineages during the current pandemic. The PANGO lineages harboring the L452R (**A and B**) and Y453F (**C and D**) and their epidemic dynamics are summarized. (**A and C**) Distribution of the L452R and Y453F mutants during the current pandemic. The top 5 PANGO lineages (https://cov-lineages.org/index.html) that harbor the L452R (**A**) and Y453F (**C**) mutations are shown. The raw data are summarized in **Table S3**. (**B and D**) Epidemic dynamics of the L452R-harboring B.1.427/429 lineage in California state, the USA (**B, top**) and the USA (**B, bottom**) and the Y453F-harboring B.1.298 lineage in Denmark (**D**). The numbers of the sequences harboring mutation per day (left y-axis, bars) and the numbers of total sequences per day (right y-axis, dots), from January 22, 2020 to March 6, 2021, are summarized. Note that a L452R variant isolated from gorilla and three Y453F variants isolated from cats are not included. See also **Figure S2** and **Tables S3 and S4**.

For the Y453F mutation, 1,274 out of the 1,380 mutated sequences belong to the B.1.1.298 lineage, which has been exclusively detected in Denmark (**Table S3**). The oldest sequence that contains the Y453F mutation in the B.1.1.298 lineage was isolated from a human in Denmark on April 20, 2020 (GISAID ID: EPI_ISL_714253) (**Figure 4C**). Intriguingly, the B.1.1.298 variants containing either Y453 or F453 are detected not only in humans but also in minks (**Figure 4D**).

The phylogenetic analysis of the whole genome sequences of the B.1.1.298 lineage SARS-CoV-2 suggested multiple SARS-CoV-2 transmissions between humans to minks (**Figure S2**). Additionally, the three sequences that contain the Y453F mutation were isolated from cats in Denmark: the two sequences out of them (GISAID ID: EPI_ISL_683164 and EPI_ISL_683166) made a single clade, while the other one (GISAID ID: EPI_ISL_683165) had a distinct origin (**Figure S2**). These results suggest that this SARS-CoV-2 variant has transmitted from humans to cats multiple times and some of them may spread among Danish cat population. However, the epidemic of a fraction of the B.1.1.298 lineage containing the Y453F mutation in Denmark peaked during October to November, 2020, and subsequently, gradually reduced (**Figure 4D**). The variant containing the Y453F mutation was last collected in Denmark on January 18, 2021 (GISAID ID: EPI_ISL_925998) and it has not been reported worldwide since then (**Figure 4D**).

## Discussion

In the present study, we demonstrated that at least two naturally occurring mutations in the SARS-CoV-2 RBM, L452R and Y453F, escape from an HLA-restricted cellular immunity and further reinforce the affinity to viral receptor ACE2. We further demonstrate that the L452R mutation increase the stability of S protein, viral infectivity and thereby enhances viral replication. Our data suggests that the L452R mutant escapes from the HLA-A24-restricted cellular immunity and further strengthens its infectivity.

Lines of recent studies have shown the emergence of the SARS-CoV-2 variants that evade the anti-SARS-CoV-2 neutralizing humoral immunity (Baum et al., 2020; Chen et al., 2021; Garcia-Beltran et al., 2021; Hoffmann et al., 2021a; Liu et al., 2021b; Liu et al., 2021c; McCarthy et al., 2021; Wang et al., 2021; Weisblum et al., 2020) and have alerted the risk of the spread of immune escape variants. In addition to humoral immunity, Zuo et al. have recently reported that functional SARS-CoV-2-specific cellular immune responses are retained at least 6 months following infection (Zuo et al., 2021). Le Bert et al. have also shown that functional cellular immune responses can contribute to controlling the disease progression of COVID-19 (Le Bert et al., 2021). These observations suggest the importance of cellular immunity eliciting efficient antiviral effects. However, the possibility of the emergence of SARS-CoV-2 variants that can evade cellular immunity has not been addressed yet. Here we demonstrated the L452R and Y453F mutations can contribute to escaping from an HLA-restricted cellular immunity. In addition to our findings, recent papers have documented that the L452R (Deng et al., 2021; Li et al., 2020) and Y453F substitutions (Baum et al., 2020; Hoffmann et al., 2021b) are potentially resistant to neutralization antibodies, suggesting that these mutants can evade both humoral and an HLA-restricted cellular immunity. Furthermore, we demonstrated that the L452R mutation significantly improve the viral replication capacity by increasing the binding affinity to human ACE2 and protein stability. Altogether, our findings suggest that the emergence of the variants that can escape from the HLA-restricted cellular immunity and further enhance viral replication capacity is another potential risk for deteriorating the COVID-19 pandemic situation.

As suggested in previous reports (Koopmans, 2021; Oude Munnink et al., 2021; WHO, 2020b), our data showed that the B.1.1.298 variant possessing the Y453F substitution is closely associated with the outbreak in minks in Denmark. Although it remains unclear whether the emergence of the Y453F mutant potentially associates with the evasion from the acquired immunity in mink, here we showed that this mutation can be resistant to the HLA-A24-resticted human cellular immunity. Because the Y453F mutation did not increase the infection efficacy using mink ACE2, our results suggest that the emergence of this mutant is not due to improving viral fitness to mink. Nevertheless, the host range of SARS-CoV-2, in terms of the use of ACE2 molecule for infection receptor, is broad in a variety of mammals (Liu et al., 2021a; OIE, 2021). More importantly, although murine ACE2 cannot be used for the infection of prototype SARS-CoV-2, recent studies have revealed that some SARS-CoV-2 variants including the B.1.351 and P.1 variants gained the ability to use murine ACE2 for infection and expanded their host range to mice (Li et al., 2021; Montagutelli et al., 2021). In addition to the evasion from human acquired immunity, zoonotic and zooanthroponosis SARS-CoV-2 transmissions can contribute to the accumulation of mutations in the spreading viruses and further impact viral phenotypes including infectivity, replication efficacy, pathogenicity, transmissibility and even host range. Therefore, the surveillance on the emergence of novel variants even in nonhuman mammals and assessing their potentials to adapt to use nonhuman ACE2 for infection receptor will be critical.

In contrast to the B.1.1.298 variant, the B.1.427/429 variant that harbors L452R substitution seem to emerge during the spread in human population, particularly in the California state in the USA, one of the hot spots of the SARS-CoV-2 outbreak in the USA [https://coronavirus.jhu.edu (as of April 2, 2021)]. Because the L452R mutation reinforces the binding affinity to human ACE2 and further enhances viral replication capacity, this variant might have emerged to improve viral fitness in humans. Another possibility is that the L452R mutant has emerged to evade the HLA-A24-restricted cellular immunity: HLA-A24 is relatively predominant in East Asian individuals (Gonzalez-Galarza et al., 2020), and the proportion of Asian American in California is highest in the USA (CDC, 2019). For instance, the proportion of the HLA-A24-positive individuals is ∼20% in San Diego (Moore et al., 2018), a city of California, where more than 270,000 SARS-CoV-2 infection cases reported so far [https://coronavirus.jhu.edu (as of April 2, 2021)]. Therefore, it might be conceivable to assume that the emergence of the L452R mutant (or the B.1.427/429 lineage) was driven by the HLA-A24-mediated cellular immunity.

In addition to the escape from antiviral acquired immunity, recent studies have shown that the emerging variants during the current pandemic, particularly the B.1.1.7 variant, can even increase viral pathogenicity and the mortality of COVID-19 (Challen et al., 2021; Davies et al., 2021; Grint et al., 2021). Importantly, the HLA-A24 individuals are relatively highly frequent in East and Southeast Asian countries, such as Japan (Japanese, allele frequency=0.364, n=1,550) and Malaysia (Malaysian, allele frequency=0.361, n=1,974) (Gonzalez-Galarza et al., 2020) (**Table S1**), where both the confirmed cases and the mortality of COVID-19 are relatively lower than European countries and the USA so far [https://coronavirus.jhu.edu (as of April 2, 2021)]. A limitation of this study is that the effects of the L452 and Y453 substitutions on viral pathogenicity, mortality and transmissibility remain unaddressed. To fully characterize the virological features of these mutants, further investigations using animal models and epidemiological data will be required. Nevertheless, here we showed direct evidence suggesting that the mutations in the RBM including L452R (in the B.1.427/429 lineage) and Y453F (in the B1.1.298 lineage) potentially escape from the HLA-A24-resticted cellular immunity, and further, the L452R mutant increase its replication capacity. Therefore, these variants, particularly those possessing the L452R mutations, such as the B.1.427/429 lineage, can be the potential threat for these countries and regions with predominant HLA-A24 individuals, and deep surveillance and tracing the epidemic of these variants will be urgently required.

## STAR★METHODS

- KEY RESOURCES TABLE
- RESOURCE AVAILABILITY

- Lead Contact
- Materials Availability
- Data and Code Availability
- EXPERIMENTAL MODEL AND SUBJECT DETAILS

- Ethics Statement
- Cell Culture
- METHOD DETAILS

- Viral Genomes and Phylogenetic Analyses
- Activation Induced Marker Assay
- Analysis of Multifunctionality and Cytotoxic Potential of CD8^+^ T cells
- Plasmid Construction
- Preparation of Soluble Human ACE2
- Preparation of the Yeast-Based SARS-CoV-2 RBD Expression System
- Analysis of the Binding Affinity of the SARS-CoV-2 S RBD Variants to Human ACE2 by Yeast Surface Display
- Pseudovirus Assay
- Lentiviral Transduction
- Protein Structure
- SARS-CoV-2 Reverse Genetics
- Plaque Assay
- SARS-CoV-2 Infection
- Real-time RT-PCR
- QUANTIFICATION AND STATISTICAL ANALYSIS

## Supplemental Information

Supplemental Information includes 2 figures and 6 tables and can be found with this article online at http://…

## Author Contributions

C.M., M.T., J.Z., T.I., A.S., T.S.T., I.N., H.N., I.K., K.U., and K.S. performed the experiments.

S.T., T.F., G.S., and Y.M. prepared experimental materials.

J.Z. and Y.K. performed structural analysis.

S.N. performed molecular phylogenetic analysis.

A.Y., N.Shimoto, Y.N., R.M., T.T., and N.Sekiya performed clinical analysis and collected clinical samples.

C.M., M.T., J.Z., T.I., A.S., S.N., T.U. and K.S. designed the experiments and interpreted the results.

K.S. wrote the original manuscript.

C.M., J.Z., T.I., A.S., S.N, and T.U. modified the manuscript. All authors reviewed and proofread the manuscript.

The Genotype to Phenotype Japan (G2P-Japan) consortium contributed to the project administration.

## Consortia

The Genotype to Phenotype Japan (G2P-Japan) consortium: Mai Fujimi, Hirotake Furihata, Haruyo Hasebe, Kazuko Kitazato, Naoko Misawa, Mikari Motomura, Akiko Oide, Sachiko Sakata, Ryo Shimizu, Mai Suganami, Miyoko Takahashi, Jiaqi Wu, Miyabishara Yokoyama, and Yuan Yue

## Acknowledgments

We would like to thank all members belonging to The Genotype to Phenotype Japan (G2P-Japan) consortium. We thank Drs. Sho Fujiwara, Kazuaki Fukushima, Masaru Tanaka and Akifumi Imamura (Tokyo Metropolitan Cancer and Infectious Diseases Center Komagome Hospital, Japan) for supporting the collection of COVID-19 convalescent samples, Dr. Mizuki Kitamatsu and Mr. Yoshiki Aritsu (Kindai University, Japan) for supporting the preparation of synthetic peptides, Drs. Hiroyuki Kishi and Hiroshi Hamana (University of Toyama, Japan) for helpful suggestion, Dr. Kenzo Tokunaga (National Institute of Infectious Diseases, Japan) for providing pC-SARS2-S, Dr. Shuetsu Fukushi (National Institute of Infectious Diseases, Japan) for providing pTargeT-human ACE2, and Dr. Masafumi Takiguchi (Kumamoto University, Japan) for providing C1R-A2402 cells. The super-computing resource was provided by Human Genome Center at The University of Tokyo and the NIG supercomputer at ROIS National Institute of Genetics.

This study was supported in part by AMED Research Program on Emerging and Re-emerging Infectious Diseases 20fk0108163 (to A.S.), 20fk0108146 (to K.S.), 19fk0108171 (to S.N. and K.S.), 20fk0108270 (to K.S.) and 20fk0108413 (to T.I., S.N. and K.S.); AMED Research Program on HIV/AIDS 20fk0410019 (to T.U. and K.S.), 20fk0410014 (to K.S.) and 21fk0410039 (to K.S.); AMED Japan Program for Infectious Diseases Research and Infrastructure 20wm0325009 (to A.S.); JST J-RAPID JPMJJR2007 (to K.S.); JST SICORP (e-ASIA) JPMJSC20U1 (to K.S.); JST CREST JPMJCR20H6 (to S.N) and JPMJCR20H4 (to K.S); JSPS KAKENHI Grant-in-Aid for Scientific Research B 18H02662 (to K.S.); JSPS KAKENHI Grant-in-Aid for Scientific Research on Innovative Areas 16H06429 (to S.N. and K.S.), 16K21723 (to S.N. and K.S.), 17H05823 (to S.N.), 17H05813 (to K.S.), 19H04843 (to S.N.) and 19H04826 (to K.S.); JSPS Fund for the Promotion of Joint International Research (Fostering Joint International Research) 18KK0447 (to K.S.); JSPS Core-to-Core Program JPJSCCB20190009 (to T.U.); JSPS Research Fellow DC1 19J20488 (to I.K.); JSPS Leading Initiative for Excellent Young Researchers (LEADER) (to T.I.); ONO Medical Research Foundation (to K.S.); Ichiro Kanehara Foundation (to K.S.); Lotte Foundation (to K.S.); Mochida Memorial Foundation for Medical and Pharmaceutical Research (to K.S.); Daiichi Sankyo Foundation of Life Science (to K.S.); Sumitomo Foundation (to K.S.); Uehara Foundation (to K.S.); Takeda Science Foundation (to C.M., T.I. and K.S.); The Tokyo Biochemical Research Foundation (to K.S.); Mitsubishi Foundation (to T.I.); Shin-Nihon Foundation of Advanced Medical Research (to T.I.); An intramural grant from Kumamoto University COVID-19 Research Projects (AMABIE) (to C.M., T.I. and T.U.); Kumamoto University International Collaborative Research Grants (to T.U.);

Intercontinental Research and Educational Platform Aiming for Eradication of HIV/AIDS (to T.I. and T.U.); and 2020 Tokai University School of Medicine Research Aid (to S.N.). T.S.T and I.N. are the recipients of the doctoral course scholarship from Japanese Government.

---

**Table S1.** Distribution of HLA-A24 allele in each population, related to Figure 1

**Table S2.** Countries where the naturally occurring mutations in the residues 448-456 of the SARS-CoV-2 S protein were isolated, related to Figure 1

**Table S3.** The PANGO lineages harboring the L452R and Y453F mutations, related to Figure 4

**Table S4.** The PANGO lineages dominantly expanding in the USA, related to Figure 4

**Table S5.** HLA-A*24:02-positive COVID-19 convalescent samples used in this study, related to Figure 1

**Table S6.** The SARS-CoV-2 genomic region encoded in each template and the primers used for the preparation of each fragment for CPER, related to Figure 3

## STAR★METHODS

### KEY RESOURCES TABLE

### RESOURCE AVAILABILITY

#### Lead Contact

Further information and requests for resources and reagents should be directed to and will be fulfilled by the Lead Contact, Kei Sato.

#### Materials Availability

All unique reagents generated in this study are listed in the Key Resources Table and available from the Lead Contact with a completed Materials Transfer Agreement.

#### Data and Code Availability

Additional Supplemental Items are available from Mendeley Data at http://…

### EXPERIMENTAL MODEL AND SUBJECT DETAILS

#### Ethics Statement

All protocols involving the human subjects recruiting at Kyushu University Hospital, Japan, National Hospital Organization Kyushu Medical Center, Japan, and Tokyo Metropolitan Cancer and Infectious Diseases Center Komagome Hospital, Japan, were reviewed and approved by the Ethics Committee for Epidemiological and General Research at the Faculty of Life Science, Kumamoto University (approval numbers 2066 and 461). All human subjects provided written informed consent.

#### Cell Culture

Human PBMCs were obtained from a total of 15 subjects harboring HLA-A*24:02 including 12 COVID-19 convalescents and 3 seronegatives (**Table S5**). The PBMCs were purified by a density gradient centrifugation using Ficoll-Paque Plus (GE Healthcare Life Sciences, cat# 17-1440-03) and stored in liquid nitrogen until further use. The C1R cells expressing HLA-A*2402 (C1R-A2402) (Karaki et al., 1993) were maintained in RPMI1640 medium (Thermo Fisher Scientific, cat# 11875101) containing 10% fetal calf serum (FCS) and 1% antibiotics (penicillin and streptomycin; PS).

HEK293 cells (a human embryonic kidney cell line; ATCC CRL-1573) and HEK293T cells (a human embryonic kidney cell line; ATCC CRL-3216) were maintained in Dulbecco’s modified Eagle’s medium (high glucose) (Wako, cat# 044-29765) containing 10% FCS and 1% PS.

Vero cells [an African green monkey (*Chlorocebus sabaeus*) kidney cell line; JCRB0111] were maintained in Eagle’s minimum essential medium (Wako, cat# 051-07615) containing 10% FCS and 1% PS.

VeroE6/TMPRSS2 cells [an African green monkey (*Chlorocebus sabaeus*) kidney cell line; JCRB1819] (Matsuyama et al., 2020) were maintained in Dulbecco’s modified Eagle’s medium (low glucose) (Wako, cat# 041-29775) containing 10% FCS, G418 (1 mg/ml; Nacalai Tesque, cat# G8168-10ML) and 1% PS.

HEK293-C34 cells, the *IFNAR1* KO HEK293 cells expressing human ACE2 and TMPRSS2 by doxycycline treatment (Torii et al., 2021), were maintained in Dulbecco’s modified Eagle’s medium (high glucose) (Sigma-Aldrich, cat# R8758-500ML) containing 10% FCS, Blasticidin (10 μg/ml; Invivogen, cat# ant-bl-1) and 1% PS.

Expi293F cells (Thermo Fisher Scientific, cat# A14527) were maintained in Expi293 expression medium (Thermo Fisher Scientific, cat# A1435101).

### METHOD DETAILS

#### Viral Genomes and Phylogenetic Analyses

All viral genome sequences and annotation information used in this study were downloaded from GISAID (https://www.gisaid.org) as of March 15, 2021 (750,243 sequences). We used the viral nucleotide sequences that do not contain any undetermined nucleotides in the region coding S protein for the analysis (581,367 sequences). The SARS-CoV-2 variants containing the L452R or Y453F mutation were sorted from the verified 581,367 sequences (**Tables S2 and S3**). To infer the phylogeny of B.1.1.298 lineage (**Figure S2**), we collected the 657 sequences belonging to the B.1.1.298 lineage that do not contain any undetermined nucleotides. We aligned whole genome sequences by using FFT-NS-2 program in an MAFFT suite v7.467 (Katoh and Standley, 2013) and removed gapped regions using trimAl v1.4.rev22 with a gappyout option (Capella-Gutierrez et al., 2009). We selected GTR+I as the best-fit nucleotide substitution model using ModelTest-NG v0.1.5 (Darriba et al., 2020). Using the model, we generated a maximum-likelihood based phylogenetic tree using RAxML-NG v1.0.0 (Kozlov et al., 2019) with a bootstrap test (n=100).

#### Activation Induced Marker Assay

The expansion of antigen-specific human CD8^+^ T cells and the analysis of the surface expression levels of activation markers, CD25 and CD137, were performed as previously described (Wolfl et al., 2007). Briefly, human PBMCs were pulsed with 1 μg/ml of the SARS-CoV-2 PepTivator peptide pools (“S overlap peptides”) (Miltenyi Biotec, cat# 130-126-700) and maintained in RPMI 1640 medium (Thermo Fisher Scientific, cat# 11875101) containing 10% FCS and 30 U/ml recombinant human IL-2 (Peprotec, cat# 200-02) for 10-14 days. The *in vitro* expanded CD8^+^ T cells (i.e., the CTL lines) were stimulated with or without the NF9 peptide (NYNYLYRLF, residues 448-456 of the SARS-CoV-2 S protein; synthesized by Scrum Inc.). After the incubation at 37°C for 1 h, the cells were washed and the surface proteins (CD3, CD8, CD14, CD19, CD25 and CD137) were stained with the antibodies listed in **Key Resources Table**. The dead cells were stained with 7-aminoactinomycin D (Biolegend, cat# 420404). After the incubation for 20 min on ice, the cells were fixed with 1% paraformaldehyde (Nacalai Tesque, cat# 09154-85) and the levels of protein surface expression were analyzed by flow cytometry using a FACS Canto II (BD Biosciences). The data obtained by flow cytometry were analyzed by FlowJo software (Tree Star).

#### Analysis of Multifunctionality and Cytotoxic Potential of CD8^+^ T cells

C1R-A2402 cells were pulsed with or without the NF9 peptide or its derivatives [the NF9-L452R peptide (NYNY<u>R</u>YRLF, L5R in NF9) and the NF9-Y453F peptide (NYNYL<u>F</u>RLF, Y6F in NF9); synthesized by Scrum Inc.] at concentrations from 0.1 to 10 nM at 37°C for 1 h. The cells were washed twice with PBS, mixed with the CTL lines generated from COVID-19 convalescents (see above) and incubated with RPMI 1640 medium (Thermo Fisher Scientific, cat# 11875101) containing 10% FCS, 5 μg/ml brefeldin A (Sigma-Aldrich, cat# B7651), 2 μM monensin (Biolegend, cat# 420701) and BV421-anti-CD107a antibody (Biolegend, cat# 420404) in a 96-well U plate at 37°C for 5 h. Then, the cells were washed and the surface proteins (CD3, CD8, CD14 and CD19) were stained with the antibodies listed in **Key Resources Table**. The dead cells were stained with 7-aminoactinomycin D (Biolegend, cat# 420404). After the incubation at 37°C for 30 min, the cells were fixed and permeabilized with Cytofix/Cytoperm Fixation/Permeabilization solution kit (BD Biosciences, cat# 554714) and the intracellular proteins (IFN-γ, TNF-α and IL-2) were stained with the antibodies listed in **Key Resources Table**. After the incubation at room temperature for 30 min, the cells were washed and the levels of protein expression were analyzed by flow cytometry using a FACS Canto II (BD Biosciences). The data obtained by flow cytometry were analyzed by FlowJo software (Tree Star).

#### Plasmid Construction

The plasmids expressing the SARS-CoV-2 S protein (pCAGGS-SARS2-S) and its mutants (pCAGGS-S-L452R, pCAGGS-SARS2-S-Y453F and pCAGGS-SARS2-S-N501Y) were generated by site-directed mutagenesis PCR using pC-SARS2-S (kindly provided by Kenzo Tokunaga) (Ozono et al., 2021) as the template and the following primers: *S* forward, 5’-TTG GGTACC ATG TTT GTG TTC CTG GTG CTG-3’; *S* reverse, 5’-GTG GCGGCCGC TCT AGA TTC AGG TGT AGT GCA GTT T-3’; *S* Y453F forward, 5’-GTG GGA GGC AAC TAC AAC TAC CTC TTC AGA-3’; and *S* L452R/Y453F reverse, 5’-GTT GTA GTT GCC TCC CAC CTT-3’; *S* N501Y forward, 5’-TCC TAT GGC TTC CAA CCA ACC TAT GGA-3’; and *S* N501Y reverse, 5’-TGGTTG GAA GCC ATA GGA TTG-3’. The resultant PCR fragment was digested with KpnI and NotI and inserted into the KpnI-NotI site of pCAGGS vector (Niwa et al., 1991).

To construct the expression plasmid for human ACE2 (GenBank: NM_021804.3) (pLV-EF1a-human ACE2-IRES-Puro), the MluI-SmaI fragment pTargeT-human ACE2 (kindly provided by Shuetsu Fukushi) (Fukushi et al., 2007) was inserted into the MluI-HpaI site of pLV-EF1a-IRES-Puro (Addgene #85132). Nucleotide sequences were determined by a DNA sequencing service (Fasmac), and the sequence data were analyzed by Sequencher v5.1 software (Gene Codes Corporation).

#### Preparation of Soluble Human ACE2

To prepare soluble human ACE2, the expression plasmid for the extracellular domain of human ACE2 (residues 18-740) based on pHL-sec (Addgene, cat# 99845) (Zahradník et al., 2021a) was transfected into Expi293F cells using ExpiFectamine 293 transfection kit (Thermo Fisher Scientific, cat# A14525) according to the manufacturer’s protocol. Three days posttransfection, the culture medium was harvested, centrifuged, and filtrated through a 0.45-µm pore size filter (Thermo Fisher Scientific, cat# 09-740-114). The filtered medium was applied on a 5-ml of HisTrap Fast Flow column (Cytiva, cat# 17-5255-01) equilibrated by phosphate buffered saline (PBS) using ÄKTA pure chromatography system (Cytiva). The column was washed with PBS and the pure human ACE2 protein (residues 18-740) was eluted using the PBS supplemented with 300 mM imidazole (pH 7.4). Using Ultracel-3 regenerated cellulose membrane (Merck, cat# UFC900324), the buffer was exchanged to PBS and the purified protein was concentrated. The purity of prepared protein was analyzed by a Tycho NT.6 system (NanoTemper).

#### Preparation of the Yeast-Based SARS-CoV-2 RBD Expression System

A labeling-free yeast surface display plasmid pJYDC1 (Addgene, cat# 162458) (Zahradník et al., 2021b) encoding the SARS-CoV-2 S RBD (residues 336-528) (pJYDC1-RBD) (Zahradník et al., 2021a) was modified by the restriction enzyme-free cloning procedure (Peleg and Unger, 2014). To prepare the plasmids with the mutated RBD, megaprimers were amplified by PCR using KAPA HiFi HotStart ReadyMix kit (Roche, cat# KK2601) and the following primers: RBD L452R forward:5’-GGA CAG CAA GGT GGG AGG CAA CTA CAA CTA CAG ATA CAG ACT GTT CAG GAA GAG CAA C-3’; RBD Y453F reverse: 5’-CTC AAA TGG TTT CAG GTT GCT CTT CCT GAA CAG TCT GAA GAG GTA GTT GTA GTT GCC TCC C-3’; RBD N501Y reverse: 5’-GTA TGG TTG GTA GCC CAC TCC ATA GGT TGG TTG GAA GCC ATA GGA TTG-3’; pCT_seq Reverse: 5’-CAT GGG AAA ACA TGT TGT TTA CGG AG-3’; and pCTCON_seq Forward: 5’-GCA GCC CCA TAA ACA CAC AGT AT-3’, according to the manufacturer’s protocol. The PCR products were integrated into pJYDC1 by integration PCR as previously described (Peleg and Unger, 2014).

#### Analysis of the Binding Affinity of the SARS-CoV-2 S RBD Variants to Human ACE2 by Yeast Surface Display

The pJYDC1-based yeast display plasmids expressing SARS-CoV-2 RBD and its mutants were transformed into yeast (*Saccharomyces cerevisiae*; strain EBY100, ATCC MYA-4941) and selected by the growth on the SD-W plates (Peleg and Unger, 2014). Single colonies were grown in 1-ml liquid SD-CAA medium (Zahradník et al., 2021a) overnight at 30°C (220 rpm) and used to inoculate the expression cultures in the 1/9 medium (Zahradník et al., 2021a) with 1 nM bilirubin (Sigma-Aldrich, cat# 14370-1G). The cells were washed with PBS-B buffer [PBS supplemented with bovine serum albumin (1 g/l)] and aliquoted in analysis solutions. The analysis solutions consist of the PBS-B buffer with the 14 different concentrations (covering the range from 100 nM to 1 pM) of the human ACE2 protein (residues 18-740) that is labeled with CF640R succinimidyl ester (Biotium, cat# 92108). The volume of the analysis solution was adjusted (1-100 ml) in order to reduce the effect of ligand depletion (Zahradník et al., 2021b). The yeasts expressing the SARS-CoV-2 S RBDs were incubated with the analysis solution overnight to allow for equilibrium. Subsequently, the yeasts were washed with the PBS-B buffer, passed through a 40-µm cell strainer (SPL Life Sciences, cat# 93040), and the binding affinity to the CF640R-labeled human ACE2 protein (residues 18-740) was analyzed using an Accuri C6 flow cytometer (BD Biosciences). The fluorescent signal was processed as previously described (Zahradník et al., 2021b) and the standard non-cooperative Hill equation was fitted by nonlinear least-squares regression using Python v3.7 (https://www.python.org).

#### Pseudovirus Assay

To prepare the pseudoviruses, the lentivirus (HIV)-based, luciferase-expressing reporter viruses that are pseudotyped with the SARS-CoV-2 S protein and its derivatives, HEK293T cells (1 × 10^6^ cells) were cotransfected with 1 μg of psPAX2-IN/HiBiT (Ozono et al., 2020), and 1 μg of pWPI-Luc2 (Ozono et al., 2020), and 500 ng of the plasmids expressing parental S or its derivatives (L452R, Y453F or N501Y) using Lipofectamine 3000 (Thermo Fisher Scientific, cat# L3000015) according to the manufacturer’s protocol. Two days posttransfection, the culture supernatants were harvested, centrifuged, and treated with 37.5 U/ml DNase I (Roche, cat# Sigma-Aldrich, cat# 11284932001) at 37°C for 30 min. The amount of the pseudoviruses prepared was quantified by HiBiT assay and the measured value was normalized to the level of HIV p24 antigen as previously described (Ozono et al., 2021; Ozono et al., 2020). The pseudoviruses prepared were stored at –80°C until use.

To prepare the target cells for pseudovirus infection, HEK293T cells (1 × 10^6^ cells) were cotransfected with 250 ng of pC-TMPRSS2 (Ozono et al., 2021) and 500 ng of pC-ACE2 (a human ACE2 expression plasmid) (Ozono et al., 2021) using Lipofectamine 2000 (Thermo Fisher Scientific, cat# 11668019) according to the manufacturer’s protocol. Two days posttransfection, the transfected cells (22,000 cells/100 μl) were seeded into 96-well plates and infected with 100 μl of the pseudoviruses prepared at 4 different doses (1, 3, 5 and 10 ng of p24 antigen). Two days postinfection, the infected cells were lysed with One-Glo luciferase assay system (Promega, cat# E6130), and the luminescent signal was measured by using a CentroXS3 plate reader (Berthhold Technologies).

#### Lentiviral Transduction

Lentiviral transduction was performed as described previously (Anderson et al., 2018; Ikeda et al., 2019). Briefly, the VSV-G-pseudotyped lenvirus vector expressing human ACE2 was generated by transfecting 2.5 μg of pLV-EF1a-human ACE2-IRES-Puro plasmid with 1.67 μg of pΔ-NRF (expressing HIV-1 *gag, pol, rev*, and *tat* genes) (Naldini et al., 1996) and 0.83 μg of pVSV-G (expressing VSV-G; Addgene, cat#138479) into 293T cells (3 × 10^6^ cells) using TransIT-LT1 (Takara, cat# MIR2300) according to the manufacturer’s protocol. Two days posttransfection, the culture supernatants were harvested, centrifuged, and the supernatants were filtered with 0.45 µm pore size filter (Millipore, cat# SLGVR33RB) and collected as the lentiviral vector. The lenvirus vectors were concentrated by centrifugation (at 22,000 × g for 2 h at 4°C) and the concentrated lentiviral vectors were inoculated into the target cells and incubated at 37°C. Two days posttransduction, the transduced cells were placed under the drug selection using the culture medium containing 1 µg/ml puromycin (Invivogen, cat# ant-pr-1). The puromycin-selected cells with relatively higher ACE2 expression were sorted by a FACS Aria II (BD Biosciences) and expanded. After the expansion, the expression level of surface ACE2 was verified by a FACS Canto II (BD Biosciences). For the staining of surface ACE2, a goat anti-ACE2 polyclonal antibody (R&D systems, cat# AF933) and an APC-conjugated donkey anti-goat IgG (R&D systems, cat# F0108) were used. A normal goat IgG (R&D systems, cat# AB-108-C) was used as the negative control of this assay.

#### Protein Structure

The 3D visualization of the SARS-CoV-2 S and human ACE2 proteins (**Figures 2A and 2F**) was generated using PyMOL v2.3 (https://pymol.org/2/) with the cocrystal structure of the SARS-CoV-2 S and human ACE2 proteins (PDB: 6M17) (Yan et al., 2020). The L452R substitution (**Figure 2F**) was prepared using UCSF Chimera v1.13 (Pettersen et al., 2004).

#### SARS-CoV-2 Reverse Genetics

Recombinant SARS-CoV-2 was generated by circular polymerase extension reaction (CPER) as previously described (Torii et al., 2021). In brief, the 9 DNA fragments encoding the partial genome of SARS-CoV-2 (strain WK-521; GISAID ID: EPI_ISL_408667) (Matsuyama et al., 2020) were prepared by PCR using PrimeSTAR GXL DNA polymerase (Takara, cat# R050A). Additionally, a linker fragment encoding hepatitis delta virus ribozyme (HDVr), bovine growth hormone (BGH) polyA signal and cytomegalovirus (CMV) promoter was prepared by PCR. The corresponding SARS-CoV-2 genomic region and the templates and the primers of this PCR are summarized in **Table S6**. The obtained 10 DNA fragments were mixed and used for the CPER (Torii et al., 2021).

To produce the recombinant SARS-CoV-2, the CPER products were transfected into HEK293-C34 cells using TransIT-LT1 (Takara, cat# MIR2300) according to the manufacturer’s protocol. One day posttransfection, the culture medium was replaced with Dulbecco’s modified Eagle’s medium (high glucose) (Sigma-Aldrich, cat# R8758-500ML) containing 2% FCS, 1% PS and doxycycline (1 μg/ml; Takara, cat# 1311N). Six days posttransfection, the culture medium was harvested, centrifuged, and the supernatants were collected as the seed virus. To remove the CPER products (i.e., SARS-CoV-2-related DNA), 1 ml of the seed virus was treated with 2 μl TURBO DNase (Thermo Fisher Scientific, cat# AM2238) and incubated at 37 °C for 1 h. The complete removal of the CPER products (i.e., SARS-CoV-2-related DNA) from the seed virus was verified by PCR. To prepare the working virus of the recombinant SARS-CoV-2 for the virological experiments (**Figure 3**), 100 μl of the seed virus was inoculated into VeroE6/TMPRSS2 cells (5,000,000 cells in T-75 flask). One hour after infection, the culture medium was replaced with Dulbecco’s modified Eagle’s medium (low glucose) (Wako, cat# 041-29775) containing 2% FCS and 1% PS. Two days postinfection, the culture medium was harvested, centrifuged, and the supernatants were collected as the working virus.

To generate the recombinant SARS-CoV-2 mutants, mutations were inserted into the fragment 8 (**Table S6**) using GENEART Site-Directed mutagenesis system (Thermo Fisher Scientific, cat# A13312) and the following primers: Fragment 8_S L452R forward, 5’-CTA AGG TTG GTG GTA ATT ATA ATT ACC GGT ATA GAT TGT TTA GGA AGT CTA ATC-3’; Fragment 8_S L452R reverse, 5’-GAT TAG ACT TCC TAA ACA ATC TAT ACC GGT AAT TAT AAT TAC CAC CAA CCT TAG-3’; Fragment 8_S Y453F forward, 5’-GGT TGG TGG TAA TTA TAA TTA CCT GTT TAG ATT GTT TAG GAA GTC TAA TCT C-3’; Fragment 8_S Y453F reverse, 5’-GAG ATT AGA CTT CCT AAA CAA TCT AAA CAG GTA ATT ATA ATT ACC ACC AAC C-3’; Fragment 8_S N501Y forward, 5’-CAA TCA TAT GGT TTC CAA CCC ACT TAT GGT GTT GGT TAC CAA CCA TAC AG-3’; and Fragment 8_S N501Y reverse, 5’-CTG TAT GGT TGG TAA CCA ACA CCA TAA GTG GGT TGG AAA CCA TAT GAT TG-3’, according to the manufacturer’s protocol. Nucleotide sequences were determined by a DNA sequencing service (Fasmac), and the sequence data were analyzed by Sequencher version 5.1 software (Gene Codes Corporation). The CPER for the preparation of SARS-CoV-2 mutants was performed using the mutated fragment 8 instead of the parental fragment 8. Subsequent experimental procedures correspond to the procedure for parental SARS-CoV-2 preparation (described above). To verify the inserted mutation in the working viruses, viral RNA was extracted using QIAamp viral RNA mini kit (Qiagen, cat# 52906). Viral RNA was reversed transcribed using SuperScript III reverse transcriptase (Thermo Fisher Scientific, cat# 18080085) according to manufacturers’ protocols. The DNA fragments including the mutations inserted were obtained by RT-PCR using PrimeSTAR GXL DNA polymerase (Takara, cat# R050A) and the following primers: WK-521 22607-22630 forward: 5’-GCA TCT GTT TAT GCT TGG AAC AGG-3’; and WK-521 23342-23367 reverse: 5’-CCT GGT GTT ATA ACA CTG ACA CCA CC-3’. Nucleotide sequences were determined as described above, and the sequence chromatograms (**Figure 3A**) were visualized using a web application Tracy (https://www.gear-genomics.com/teal/) (Rausch et al., 2020).

#### Plaque Assay

One day prior to infection, 200,000 Vero cells were seeded into the 12-well plate. The virus was diluted with serum-free virus dilution buffer [1 × minimum essential medium (Temin’s modification) (Thermo Fisher Scientific, cat# 11935046) with 20 mM Hepes, non-essential amino acid (Thermo Fisher Scientifc, cat# 11140-050) and antibiotics]. After removing the culture media, Vero cells were infected with 500 µl of the diluted virus at 37 °C. Two hours postinfection, 1 ml of mounting solution [1 × minimum essential medium containing 3% FCS and 1.5% carboxymethyl cellulose (Sigma, cat# C9481-500G)] was overlaid and incubated at 37 °C. Three days postinfection, the culture media were removed, and the cells were washed with PBS three times and fixed with 10% formaldehyde (Nacalai Tesque, cat# 37152-51) or 4% paraformaldehyde (Nacalai Tesque, cat# 09154-85). The fixed cells were washed with city water, dried up, and stained with staining solution [2% crystal violet (Nacalai Tesque, cat# 09804-52) or 0.1% methylene blue (Nacalai Tesque, cat# 22412-14) in water]. The stained cells were washed with city water, dried up, and the number of plaques was counted to calculate plaque forming unit (pfu).

#### SARS-CoV-2 Infection

One day prior to infection, 10,000 cells of VeroE6/TMPRSS2 and 293-ACE2 cells were seeded into the 96-well plate. Recombinant SARS-CoV-2 (100 pfu) was inoculated and incubated at 37 °C for 1 h. The infected cells were washed and replaced with 180 µl of culture media. The culture supernatant (10 µl) was harvested at 0, 6, 24, 48 and 72 hours postinfection and used for real-time PCR to quantify the copy number of viral RNA.

#### Real-time RT-PCR

The amount of viral RNA in the culture supernatant was quantified by real-time RT-PCR as previously described (Shema Mugisha et al., 2020) with some modifications. In brief, 5 μl of culture supernatants was mixed with 5 μl of 2 × RNA lysis buffer [2% Triton X-100, 50 mM KCl, 100 mM Tris-HCl (pH 7.4), 40% glycerol, 0.8 U/μl recombinant RNase inhibitor (Takara, cat# 2313B)] and incubated at room temperature for 10 min. RNase-free water (90 μl) was added and the diluted sample (2.5 μl) was used as the template of real-time RT-PCR. Real-time RT-PCR was performed according to the manufacturer’s protocol using the One Step TB Green PrimeScript PLUS RT-PCR kit (Takara, cat# RR096A) and the following primers: Forward *N*, 5’-AGC CTC TTC TCG TTC CTC ATC AC-3’; and Reverse *N*, 5’-CCG CCA TTG CCA GCC ATT C-3’. The copy number of viral RNA was standardized by SARS-CoV-2 direct detection RT-qPCR kit (Takara, cat# RC300A). The fluorescent signal was acquired on a QuantStudio 3 Real-Time PCR systems (Thermo Fisher Scientific), a CFX Connect Real-Time PCR Detection system (Bio-Rad) or a 7500 Real Time PCR System (Applied Biosystems) was used.

### QUANTIFICATION AND STATISTICAL ANALYSIS

Data analyses were performed using Prism 7 (GraphPad Software). Unless otherwise stated, the data are presented as means with SD. Statistically significant differences were determined as described in the figure legend. Statistical details can be found directly in the corresponding figure legends.

## KEY RESOURCES TABLE

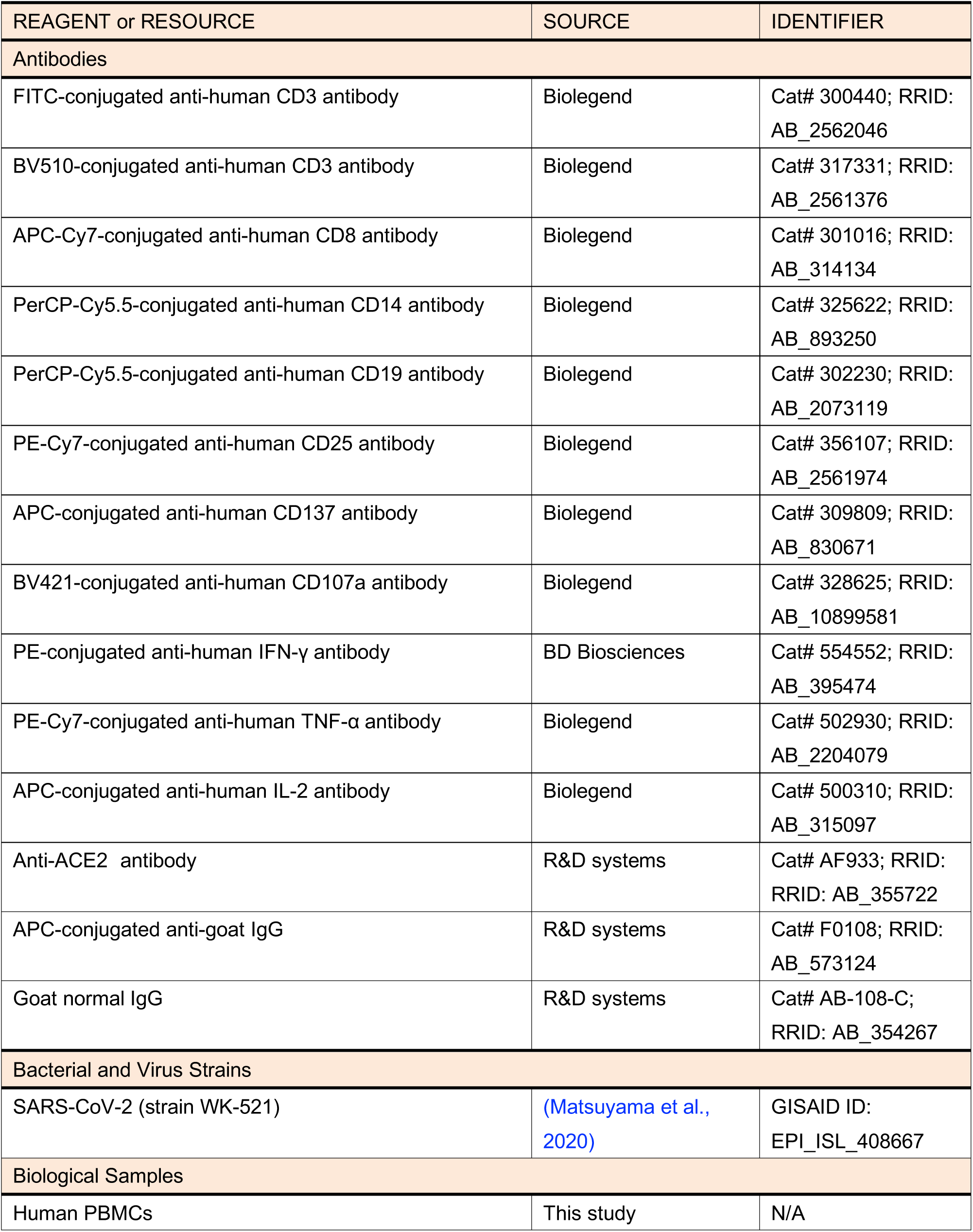
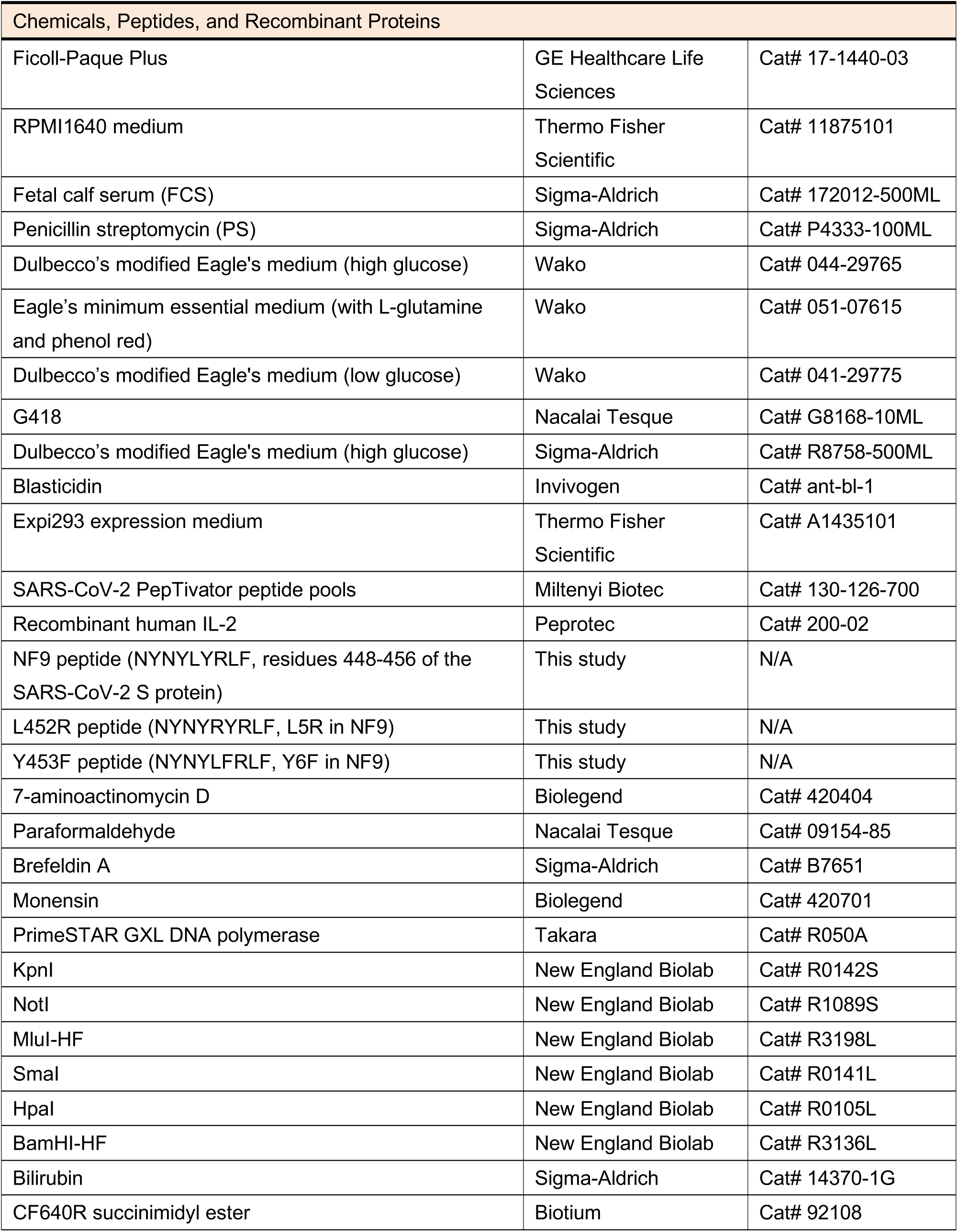
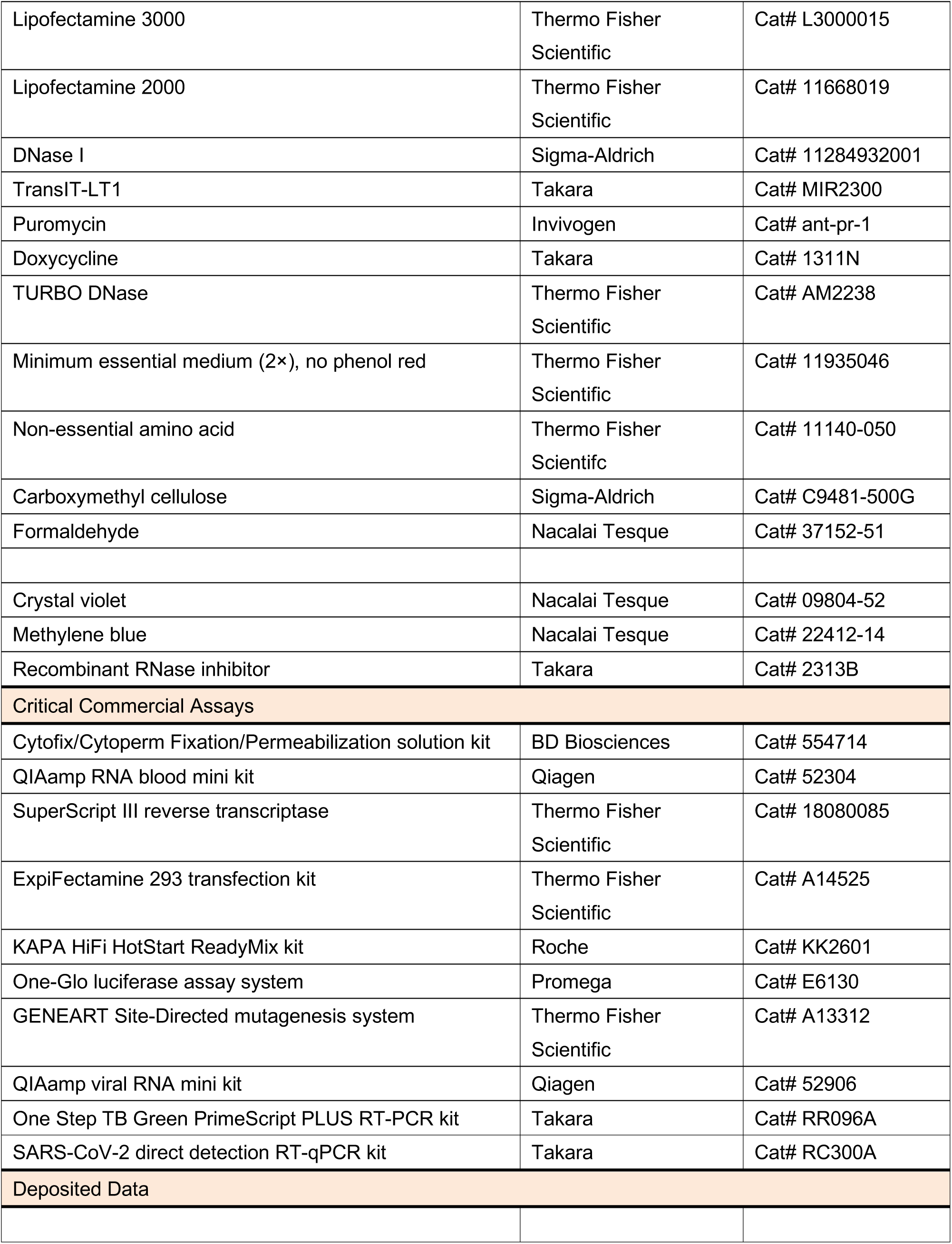
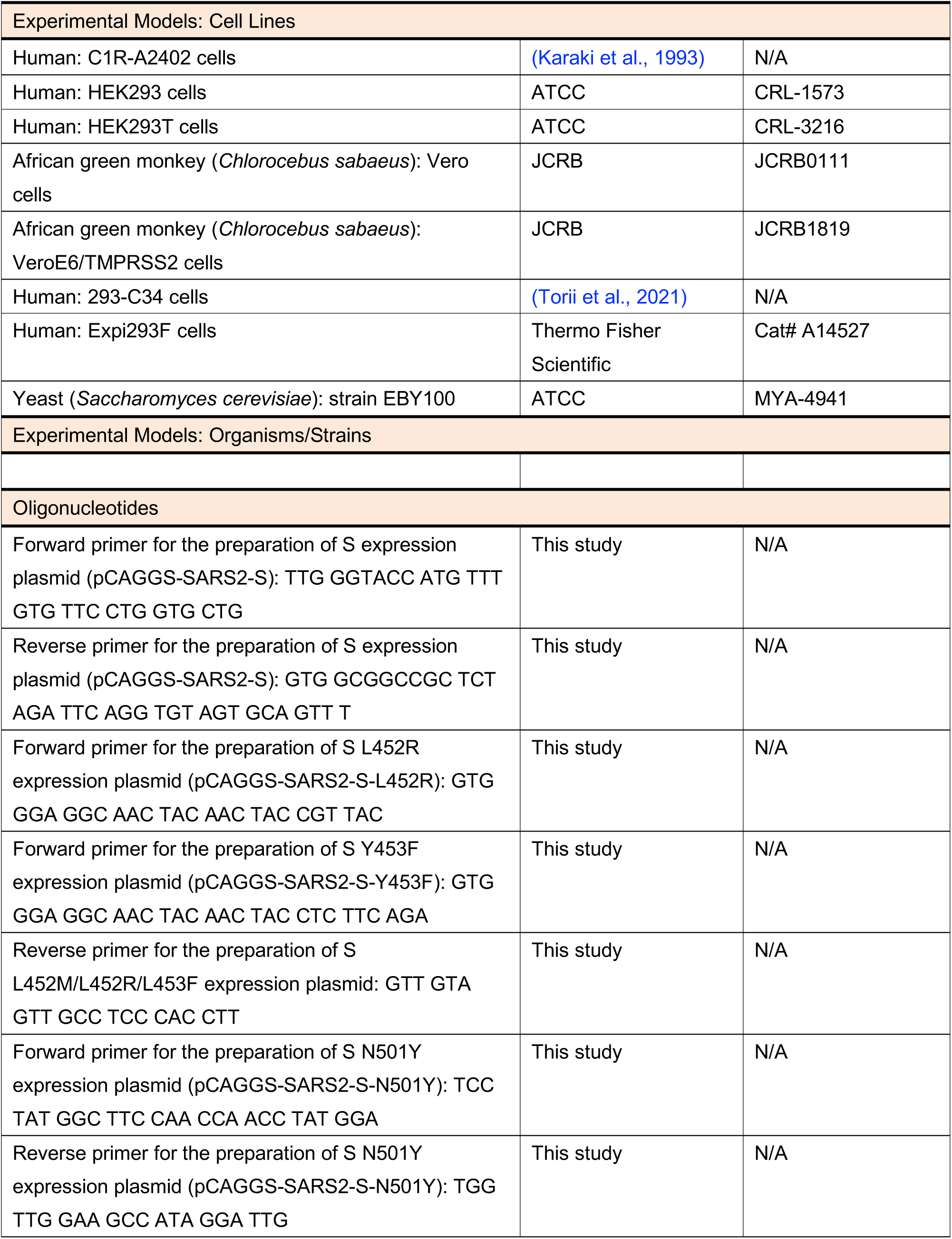
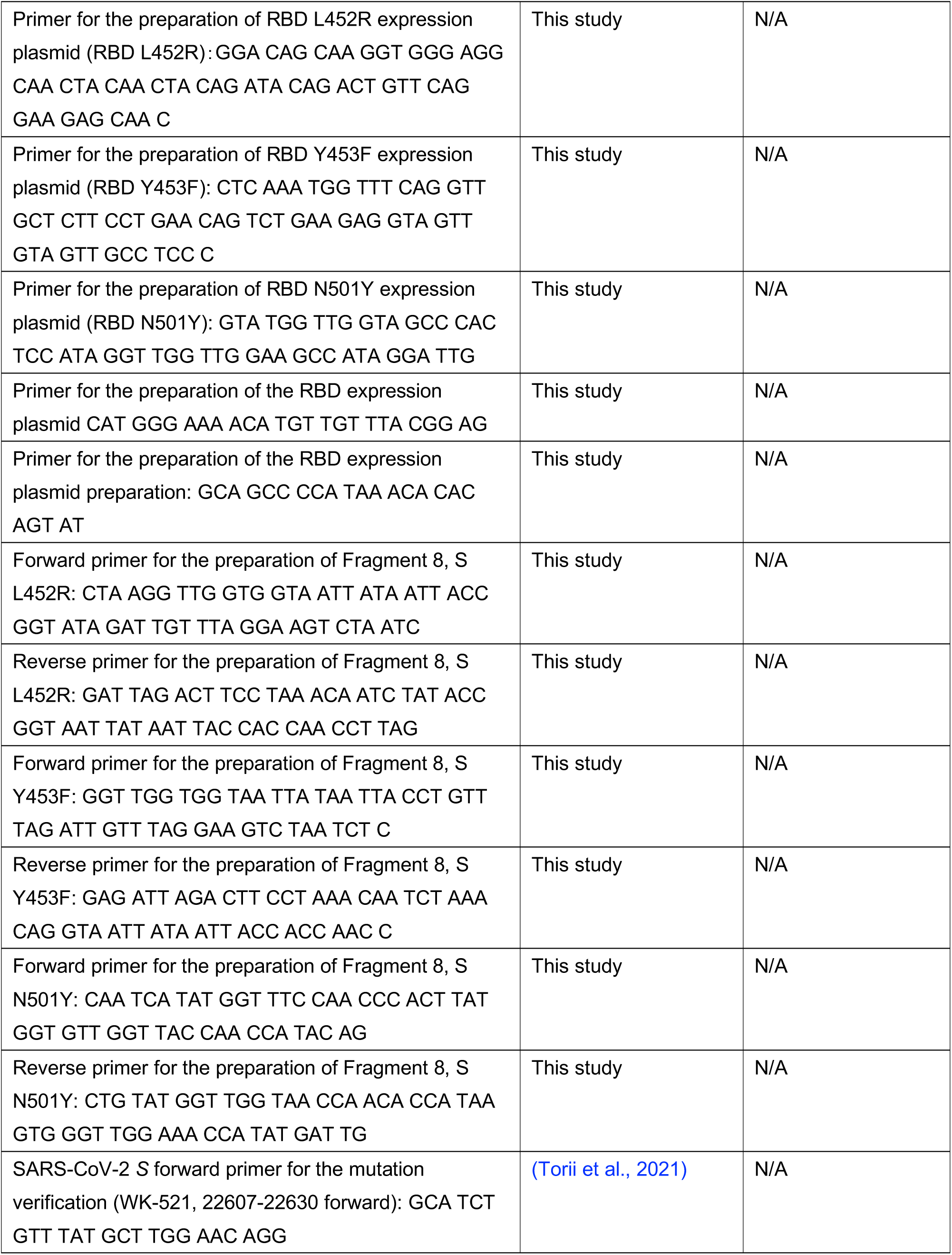
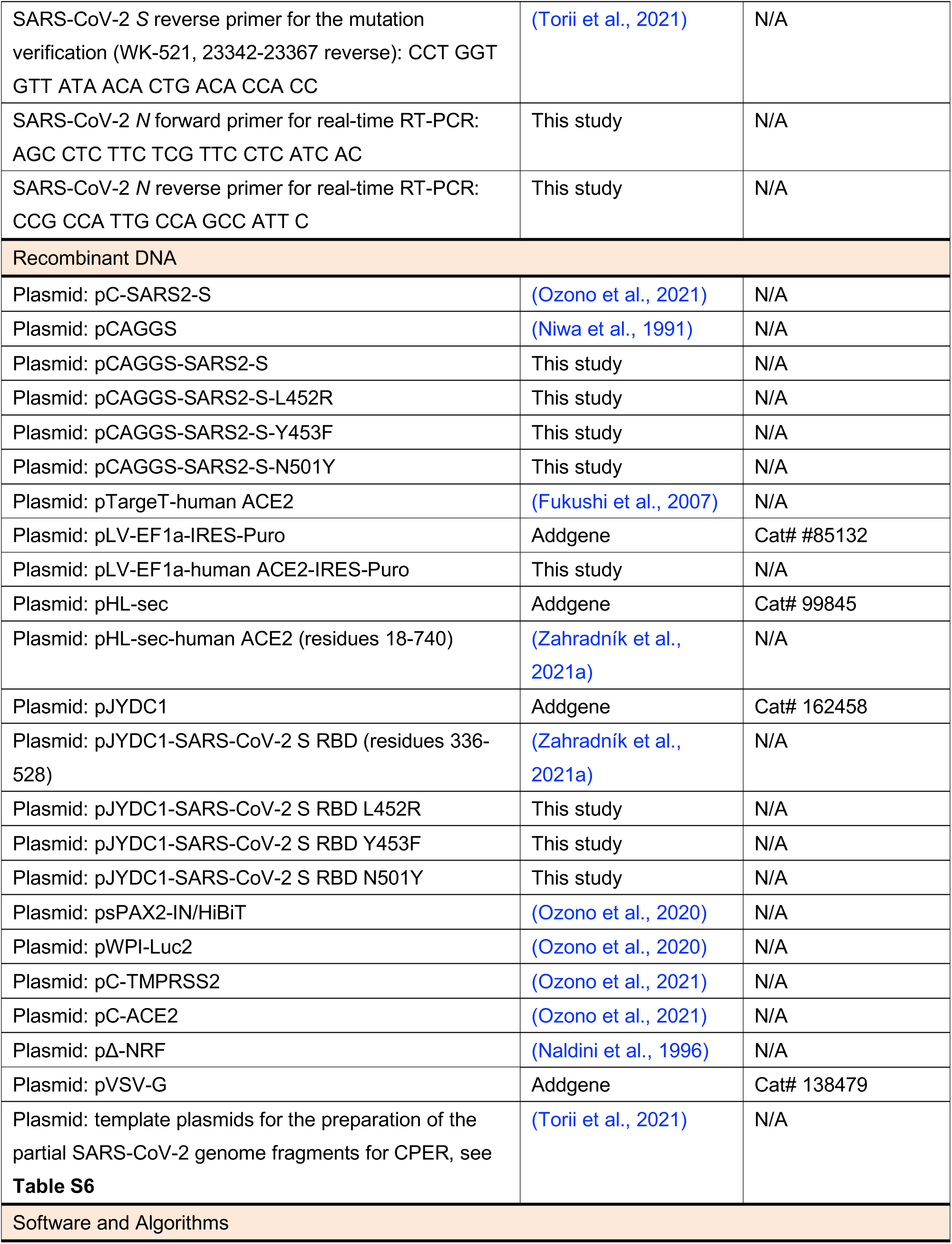
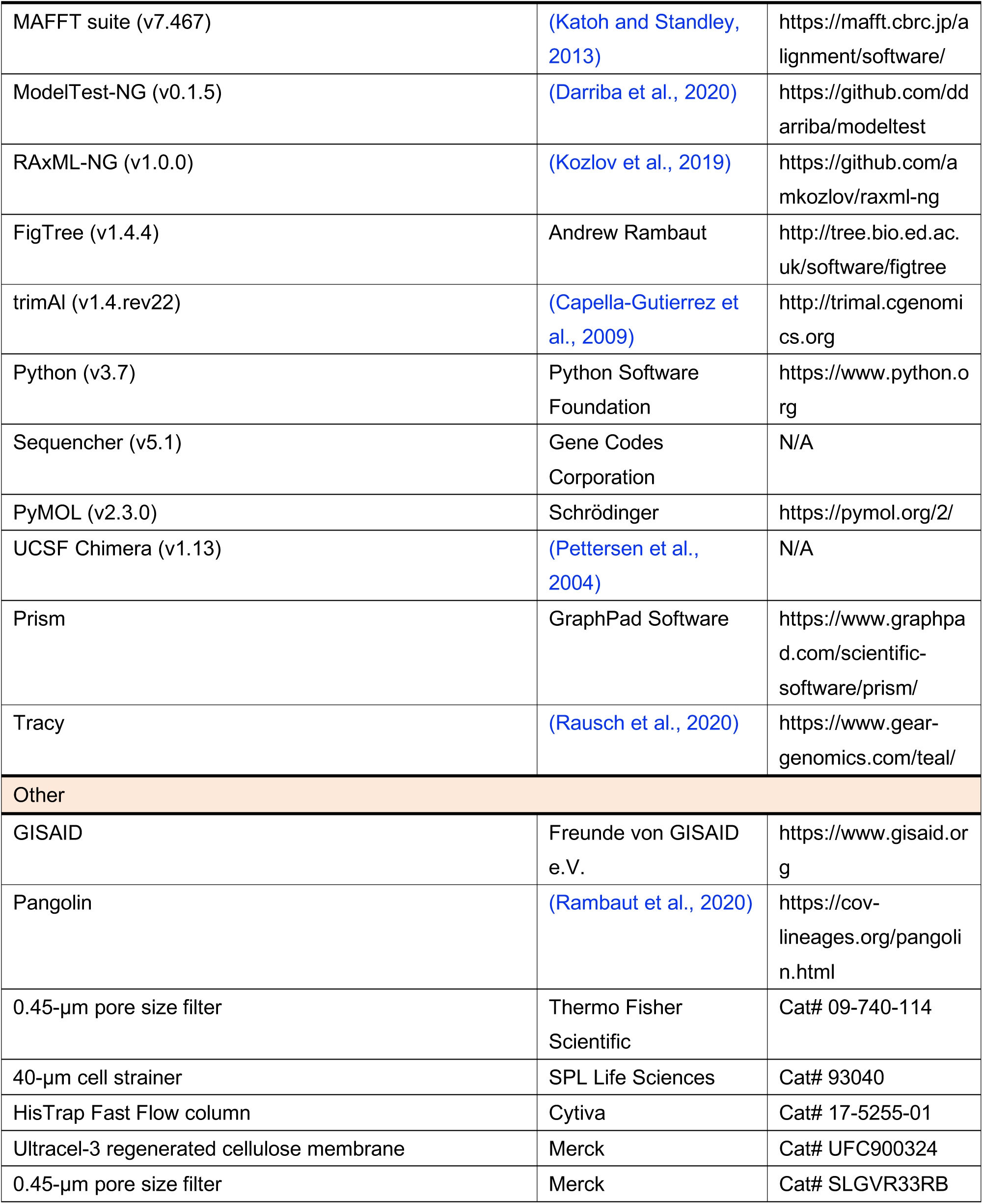

