## Supplementary material for "An emerging SARS-CoV-2 mutant evading cellular immunity and increasing viral infectivity": Figure S1 and S2

Supplementary Figures S1-S2
Supplementary Tables S2-S6

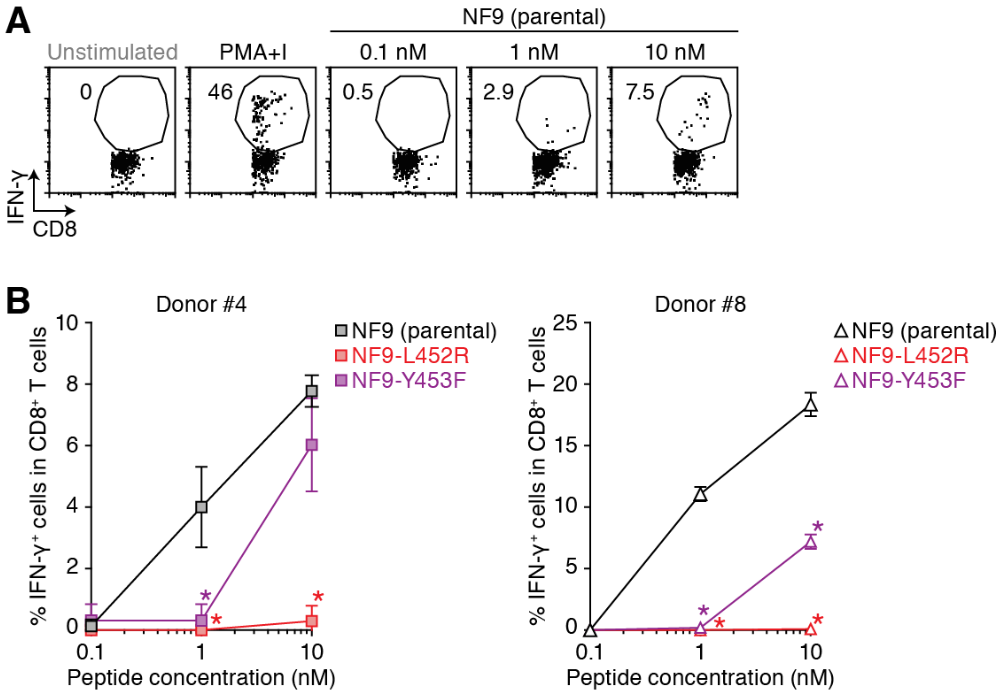


**Figure S1. IFN-γ production of HLA-A24-restricted NF9-specific CD8⁺ T cells by parental NF9 but not by its derivatives (Related to Figure 1).**

HLA-A24-restricted NF9-specific CD8⁺ T cells were stimulated with the C1R-A2402 cells pulsed with 0.1, 1 and 10 nM of NF9 peptide or PMA (10 ng/ml) and Ionomycin (1 μg/ml) (PMA+I).

(**A**) Representative plots of donor #4. The numbers in the plots indicate the percentage of IFN-γ⁺ cells in CD8⁺ T cells.

(**B**) The sensitivity of HLA-A24-restricted NF9-specific CD8⁺ T cells towards parental NF9 peptide and its derivatives (NF9-L452R and NF9-Y453F). The assay was performed in triplicate, and the means with SDs are shown. In **B**, statistically significant differences (*, *P*<0.05) versus parental NF9 at the same dose are determined by Student's *t* test.

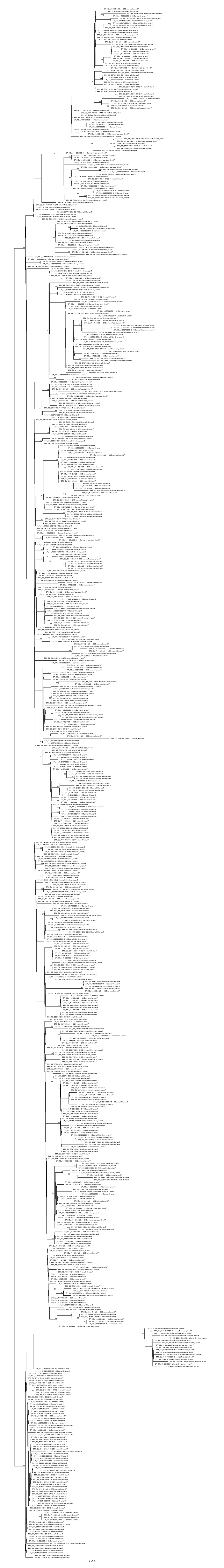

**Figure S2. A maximum-likelihood based phylogenetic tree of the 657 SARS-CoV-2 sequences that belongs to the B.1.1.298 lineage (Related to Figure 4).** GISAID ID, sampling date, country, host and amino acid at the position 453 of Spike protein were noted in each terminal node. Bootstrap values were shown in each internal node if the value was ≥50 (n=100).
